# Seascape genomics as a new tool to empower coral reef conservation strategies: an example on north-western Pacific *Acropora digitifera*

**DOI:** 10.1101/588228

**Authors:** Oliver Selmoni, Estelle Rochat, Gael Lecellier, Veronique Berteaux-Lecellier, Stéphane Joost

## Abstract

Coral reefs are suffering a major decline due to the environmental constraints imposed by climate change. Over the last 20 years, three major coral bleaching events occurred in concomitance of anomalous heat waves, provoking a severe loss of coral cover worldwide. The conservation strategies for preserving reefs, as they are implemented now, cannot cope with global climatic shifts. Consequently, researchers are advocating the set-up of preservation frameworks to reinforce coral adaptive potential. However, the main obstacle to this implementation is that studies on coral adaption are usually hard to generalize at the scale of a reef system.

Here, we study the relationships between frequencies of genetic markers with that of environmental characteristics of the sea (seascape genomics), in combination with connectivity analysis, to investigate the adaptive potential of a flagship coral species of the Ryukyu Archipelago (Japan). By associating genotype frequencies with descriptors of historical environmental conditions, we discovered six genomic regions hosting polymorphisms that might promote resistance against thermal stress. Remarkably, annotations of genes in these regions were consistent with molecular roles associated with heat responses. Furthermore, we combined information on genetic and spatial distances between reefs to predict connectivity at a regional scale.

The combination between the results of these analyses portrayed the adaptive potential of this population: we were able to identify reefs carrying potential adaptive genotypes and to understand how they disperse to neighbouring reefs. This information was summarized by objective, quantifiable, and mappable indices covering the whole region, which can be extremely useful for future prioritization of reefs in conservation planning. This framework is transferable to any coral species on any reef system, and therefore represents a valuable tool for empowering preservation efforts dedicated to the protection of coral reef in warming oceans.

## Introduction

Coral reefs are suffering a severe decline due to the effects of climate change (Hughes et al., 2017). Loss of reef is already showing catastrophic consequences for marine wildlife who depend on these structures (Pratchett et al., 2018), with disastrous aftermaths expected for human economies as well (Moberg & Folke, 1999). One of the major threats to the persistence of these ecosystems is coral bleaching (Bellwood et al., 2004): a physiological response induced by environmental stress that provokes hard-skeleton corals, the cornerstone of reefs, to separate from the symbiotic microbial algae essential for its survival (Mydlarz et al., 2010). Over the last 20 years, episodes ofcoral bleaching struck world-wide and resulted in a local coral cover loss of up to 50% (Hughes et al., 2017, 2018). Thermal stress is considered the main driver of coral bleaching (Hughes et al., 2017), but additional causes of anthropogenic origin were also identified (e.g. ocean acidification, water eutrophication, sedimentation and overfishing; Anthony et al., 2008; Ateweberhan et al., 2013; Maina et al., 2008).

Conservation efforts to mitigate the threat of coral bleaching tend to focus on restoring reefs that have undergone severe losses, as well as try to limit the impact of future bleaching events (Baums, 2008; Bellwood et al., 2004; Young et al., 2012). To achieve these aims, two main strategies are currently used: establish marine protected areas (MPAs) at reefs, and develop coral nurseries (Baums, 2008; Bellwood et al., 2004; Young et al., 2012). MPAs are designated zones in which human access and activities are severely restricted in order to alleviate the effects of local anthropogenic stressors (Lester et al., 2009). Coral nurseries are underwater gardens of transplanted colonies that can then be transplanted to restore damaged reefs (Baums, 2008; Young et al., 2012). For both conservation strategies, researchers advocate the use of methods that account for demographic connectivity such that the location of a conservation measure can also promote resistance and resilience for neighbouring sites (Baums, 2008; Krueck et al., 2017; Lukoschek et al., 2016; Palumbi, 2003; Shanks et al., 2003). Despite the observed beneficial effects of these conservation policies worldwide (Cinner et al., 2016; Rodgers et al., 2017; Selig & Bruno, 2010), these solutions do not confer resistance against the temperature oscillations associated with the last mass bleaching events (Baums, 2008; Hughes et al., 2017). Coral reefs that had experienced previous thermal stress were found to be more resistant to subsequent heat waves (Hughes et al., 2019; Krueger et al., 2017; Penin et al., 2013; Thompson & van Woesik, 2009), but to date this information is neglected in conservation actions (Baums, 2008; Maina et al., 2011). There is an urgent need to understand whether these observations are due to evolutionary processes and, if so, to determine how the underlying adaptive potential could be included in predictions of climate change responses and in conservation programs (Baums, 2008; Logan et al., 2014; Maina et al., 2011; van Oppen et al., 2015a).

To this end, seascape genomics tools are likely to play an important role. Seascape genomics is the marine counterpart of landscape genomics, a branch of population genomics that investigates adaptive potential through field-based experiments (Balkenhol et al., 2017). Samples that are collected across a landscape are genotyped using next-generation-sequencing techniques, providing thousands of genetic variants, while simultaneously the environmental variables of the study area are characterized, usually using remote-sensing data to describe specific local climatic conditions (Leempoel et al., 2017). Genomics and environmental information are then combined to detect genetic polymorphisms associated with particular conditions (*i*.*e*., potentially adaptive genotypes; Rellstab et al., 2015). This approach has been applied to many terrestrial species, and is increasingly being used to analyse marine systems in what is referred to as *seascape genomics* (exhaustively reviewed in Riginos, Crandall, Liggins, Bongaerts, & Treml, 2016). To our knowledge no seascape genomics experiment has yet been applied to reef corals. In fact, adaptation of these species has been mostly studied via transplantation assays coupled with aquarium conditioning, which is a time- and resource-4 demanding approach that is often restricted to a couple of reefs experiencing contrasting conditions (Howells et al., 2013; Krueger et al., 2017; Palumbi et al., 2014; Sampayo et al., 2016; Ziegler et al., 2017). Genotype-environment associations studies have also been conducted on corals, but have used either a limited number of markers (<10 SNPs in Lundgren, Vera, Peplow, Manel, & van Oppen, 2013), a restricted number of locations (two in Bay & Palumbi, 2014), or focused on populations with restricted gene flow (L. Thomas et al., 2017). Contrary to these previous studies, a seascape genomics approach should cover ecologically meaningful spatial scales and be able to distinguish the pressures caused from different climatic conditions, as well as account for confounding effects of demographic processes (Balkenhol et al., 2017).

In the present study, we applied a seascape genomics framework to detect coral reefs that are carrying potentially adaptive genotypes, and in turn, to show how conservation policies could implement the results. Our study focuses on *Acropora digitifera* of the Ryukyu Archipelago in Japan (Figure 1), an emblematic species of the Indo-Pacific and flagship organism for studies on corals genomics (Shinzato et al., 2011). We first analysed the convergence between genomic and environmental information to i) detect loci potentially conferring a selective advantage, and ii) develop a model describing connectivity patterns. Next, we took advantage of these findings to indicate which reefs were more likely to be carrying adaptive genotypes and to evaluate their interconnectedness with the rest of the reef system. Finally, we propose an approach to implement the results obtained into conservation planning. Overall, our work provides tools for the interface between conservation genomics and marine environmental sciences, which are likely to empower preservation strategies for coral reefs into the future.

**Figure 1.**
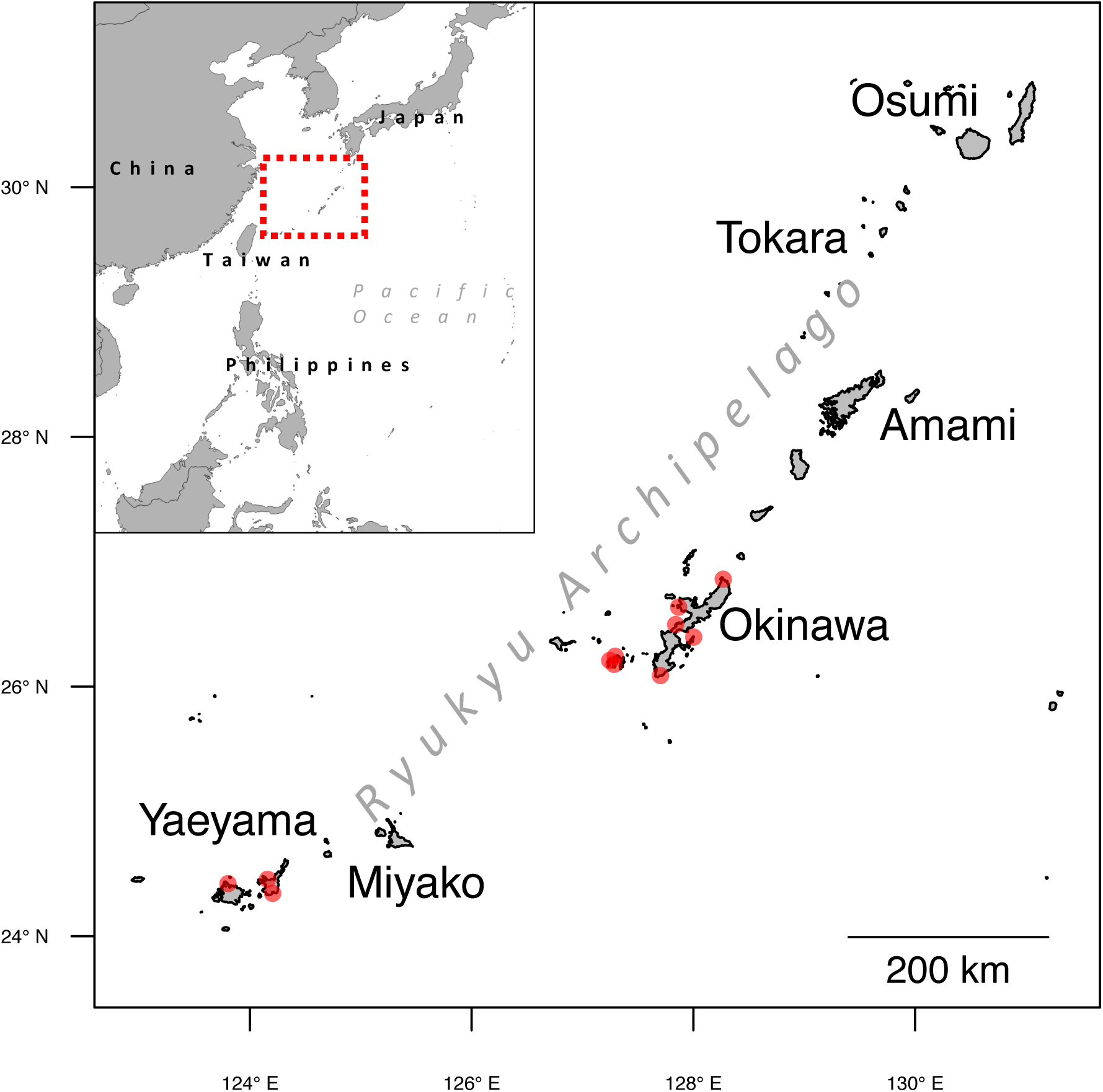
The study area. The Ryukyu Archipelago extends for more than 1,000 km in the north-western Pacific Ocean. The red circles display the 11 sites where samples were collected for the seascape genomics analysis (adapted from Shinzato, Mungpakdee, Arakaki, & Satoh, 2015).

## Materials and methods

Our framework is structured on two axes of analysis and prediction: one focusing on the presence of adaptive genotypes (seascape genomics), and the other on population connectivity (Fig. 2). The seascape genomics analysis (Fig. 2A) combines genomic data with environmental information to uncover potentially adaptive genotypes at sampling sites. The model describing this relationship is then used to predict, at the scale of the whole study area, the probability of the presence of adaptive genotypes (Fig. 2B). In the connectivity study (Fig. 2C), we made a model describing how distances based on sea currents (calculated on the basis of remote sensing data) correspond to the genetic separation between sites. This model is then used to predict connectivity of sites at the study area scale (Fig. 2D). Finally, the predictions of where the adaptive genotypes are more likely to exist, and of how the reef system is interconnected, allow the assessment of adaptive potential across the whole study area (Fig. 2E).

**Figure 2.**
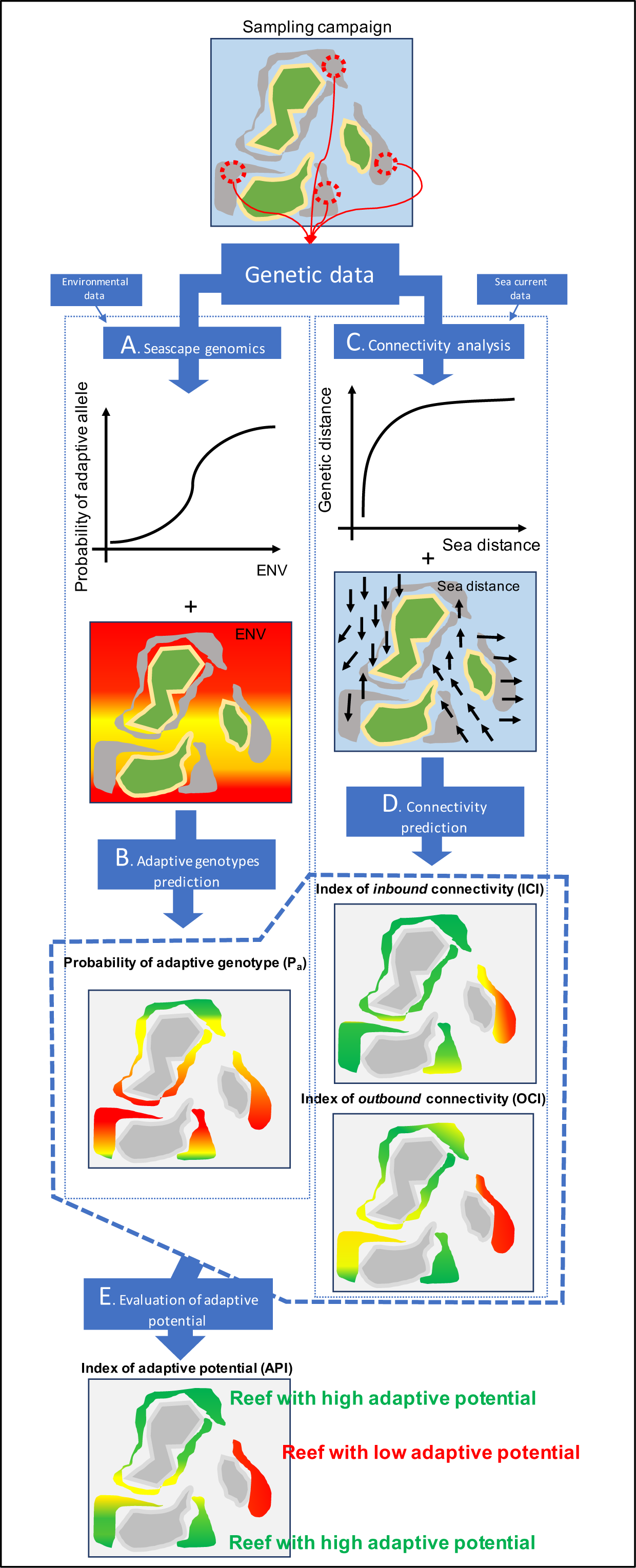
Workflow of the methods. The starting point for the analysis is the generation of genetic data describing the genotypes observed at different sampling locations (in this example, 4 sampling sites). In the seascape genomics analysis (A), these data are combined with environmental information to uncover genotypes whose frequencies are associated with specific climatic conditions (ENV). Such genotypes are defined as potentially adaptive. The model describing this link is then applied to environmental data at the scale of the whole reef system (B), to predict the probability of presence of the adaptive genotypes (green: high probability; red: low probability). The genetic data are also combined with sea current information to build a connectivity model (C) describing how sea distances correspond to genetic separation between sampling sites. This model is fitted with sea distance between all the reefs of the study area to predict (D) patterns of connectivity from (outbound) and to (inbound) each reef (green: high connectivity; red: low connectivity). Finally, predictions of the presence of adaptive genotypes and connectivity patterns are combined to assess the adaptive potential across the study area (E): reefs that are connected with sites that are likely to carry the adaptive genotype will have a higher adaptive potential (green), while those that are isolated will have lower adaptive potential (red).

### Genomic dataset

The genomic data used come from a publicly available dataset consisting of 155 geo-referenced colonies of *A. digitifera* from 12 sampling locations (13±5 colonies per site) of the Ryukyu Archipelago in Japan (Fig. 1; Bioproject Accession PRJDB4188). These samples were sequenced using a Whole-Genome Sequencing approach in the scope of a population genomics study. Details on how samples were collected and processed for genomic analysis can be found in Shinzato et al. (2015).

Genomic data were processed using the Genome Analysis Toolkit framework (GATK; McKenna et al., 2010) following the recommended pipeline (the “GATK Best Practices”; Van der Auwera et al., 2013) with the necessary modifications for coping with the absence of reliable databases of known variants for this species. In short, the *A. digitifera* reference genome (v. 1.1, GenBank accession: GCA_000222465.2; Shinzato et al., 2011) was indexed using bwa (v. 0.7.5a, Li & Durbin, 2009), samtools (v. 1.9, Heng Li et al., 2009) and picard-tools (v. 1.95, http://broadinstitute.github.io/picard) and raw sequencing reads were aligned using the bwa *mem* algorithm. The resulting alignments were sorted, marked for duplicate reads, modified for read-group headers and indexed using picard-tools. Next, each alignment underwent an independent variant discovery using the GATK HaplotypeCaller tool (using the ERC mode and setting the --minPruning flag to 10) and genotypes were then jointly called by the GATK GenotypeGVCFs tool in random batches of 18 samples to match our computational power (18 CPUs). The variant-calling matrices of the different batches were then joined and filtered in order to keep only bi-allelic Single Nucleotide Polymorphisms (SNPs) using the GATK CombineVariants and SelectVariants tools, respectively. This resulted in a raw genotype matrix counting ∼1.2 M of SNPs. Subsequently, we used the GATK VariantAnnotator tool to annotate variants for Quality-by-depth and filtered for this value (<2), read coverage (>5 and <100 within a sample), minor allele frequency (>0.05), major genotype frequency (<0.95) and missing rate of both individuals and SNPs (<0.1) using the GATK VariantFiltrationTool and custom scripts in the R environment (v. 3.5.1, R Core Team, 2016). Finally, we filtered for linkage disequilibrium using the *snpgdsLDpruning* function of the SNPrelate R package (v. 1.16, LD threshold=0.3; Zheng et al., 2012). This pipeline produced the filtered genotype matrix consisting of 136 individuals and 7,607 SNPs.

Natural hybridization and transient species boundaries have been observed in *Acropora* species (Van Oppen et al., 2002). We investigated these hypotheses by running a preliminary analysis of fixation index (Fst) variation by genomic position using the R KRIS package (v. 1.1; Chaichoompu et al., 2018). Since we found no genomic islands of low-recombination (i.e. high Fst; Nosil et al., 2009) between the populations of Kerama, Yaeayama and Okinawa (Fig. S1) we excluded the possibility of presence of genetically isolated groups in the dataset. Importantly, previous studies on this coral population did not report hybridization with other species, neither the presence of cryptic species nor isolated sub-populations (Nakajima et al., 2010; Nishikawa, 2008; Shinzato et al., 2015).

### Environmental data

Twelve georeferenced datasets describing atmospheric and seawater conditions were retrieved from publicly available resources (EU Copernicus Marine Service, 2017; NASA, 2016; National Oceanic and Atmospheric Administration, 2017; Table S1). These remote sensing derived datasets provide monthly or daily variables measured over several (on average 14) years before the genetic data were sampled (2010; Shinzato et al., 2015). The data cover the entire study area (Fig. 1), with a spatial resolution ranging from 25 km to 4 km (Tab. S1). We processed these variables in the R environment using the *raster* package (v. 2.8, Hijmans, 2016) to compute monthly averages and standard deviations for the entire study period. Furthermore, Sea Surface Temperature (SST) measurements were used to compute a Degree Heat Week (DHW) frequency index, representing the percentage of days during which the bi-weekly accumulated heat stress exceeded 4 °C (Liu et al., 2003; Logan et al., 2014). SST and sea surface salinity records were combined to produce estimates of seawater pH (Covington & Whitfield, 1988), dissolved inorganic carbon (Loukos et al., 2000), and alkalinity (Lee et al., 2006). Bathymetry data (Ryan et al., 2009) were used to retrieve the depth at sampling locations. Finally, population density data (CIESIN Columbia University, 2010) were averaged in a 50 km buffer area to produce a surrogate-variable for anthropogenic pressure (Welle et al., 2017).

We used the geographic coordinates associated with each sample to characterize the environmental conditions using the QGIS point sampling tool (v. 2.18.25, QGIS development team, 2009). For the predictive step of our study (Fig. 2C) at the scale of the whole reef system we retrieved the shapes of the reefs of the region (UNEP-WCMC et al., 2010) and reported them into a regular grid (cell size of 5×5 km) using QGIS. For the reef cells smaller than a 5- by-5 km square, we calculated the actual area (in km^2^). Reefs from the neighbouring regions (Taiwan and Philippines, Fig. 1) were also included to avoid border-effects in computations. Environmental conditions were assigned to each reef-cell using the average function of the QGIS zonal statistics tool.

### Seascape genomics

We performed the genotype-environment association analysis using the logistic regression method implemented within the SamBada software (v. 0.7; Stucki et al., 2017), customized to speed-up multivariate models computation in the Python environment (v. 2.7; Python Software Foundation, 2018) using the *pandas* (v. 0.23.4; McKinney, 2010) and *statsmodels* (v. 0.9; Seabold & Perktold, 2010) libraries. The SamBada approach allows proxy variables of genetic structure to be included in the analysis in order to avoid possible confounding effects (patterns of neutral genetic variation mimicking signals of adaptation to the local environment; Holderegger et al., 2008). Here we performed a discriminant analysis of principal components (DAPC) on the SNPs genotype matrix using the R package *adegenet* (v. 2.1.1; Jombart, 2008). This procedure highlighted a main separation between two groups of samples along the first discriminant function (Fig. S2). The latter was therefore used as co-variable in association models. We then ran the pipeline for the discovery of adaptive signals described below and summarized in a schematic in Fig. S3. In order to speed up calculations, the 315 environmental variables were grouped into 29 clusters of highly correlated (Pearson method, |R|>0.7) descriptors in the R environment (Fig. S3A). For each of these groups, one variable was randomly selected to build logistic models against the three genotypes of each SNP (Fig. S3B). In total, the SamBada instance evaluated 661,809 association models (29 environmental groups against 3 x 7,607 SNPs; Fig. S3C, S2D) that were subsequently analysed in the R environment. For each association-model related to the same environmental variable, p-values of G-scores (G) and Wald-scores (W) were corrected for multiple testing using the R *q-value* package (v. 9 2.14, Storey, 2003). Association models scoring q<0.001 for both statistics were deemed significant (Fig. S3E). If a SNP was found in more than one significant association, only the best model (according to the value of G) was kept. For each significant association retained, we then calculated all the models built with the focal SNP against the other environmental variables from the same correlated descriptor cluster, and we looked for the best association model (according to G; Fig. S3F). The best association model for each significant SNP is hereafter referred to as the significant genotype-environment association (SGEA). Finally, we visualized the logistic regression of each significant SGEA using the R *popbio* library (v. 2.4.4; Stubben & Milligan, 2007).

### Annotation of seascape genomics results

We then annotated the genomic neighbourhood of each SGEA in order to discover genes potentially linked to an adaptive role. We set the size of the search window to ±250 kbs around the concerned SNP of each SGEA. This window size was selected because genes(s) possibly linked to a mutation may lay up to hundreds of kbs away (Brodie et al., 2016; Visel et al., 2009), and this window size corresponds approximately to the scaffold N50 statistics of the reference genome (*i*.*e*. half of the genome is contained within scaffolds of this size or longer). For every SGEA, the annotation procedure was performed as follows. Based on the NCBI annotation of the reference genome (https://www.ncbi.nlm.nih.gov/genome/annotation_euk/Acropora_digitifera/100/), we retrieved all the predicted genes falling within the ±250 kbs window. Next, we retrieved the predicted protein sequences related to these genes and ran a similarity search (blastp, (Madden & Coulouris, 2008) against metazoan protein sequences in the swissprot database (release 2019_07; Boeckmann et al., 2003). For every predicted gene, only the best significant match (E-score threshold < 10^−7^) was kept. Finally, predicted genes were annotated with the eukaryotic cluster of orthologous genes (KOG; Jensen et al., 2008) annotation from the matching swissprot entry. For every KOG we calculated the relative frequency across the *A. digitifera* genome. This was obtained by dividing the genome into 500 kbs windows and by calculating the percentage of windows in which the KOG was observed.

### Probability of Presence of Adaptive Genotypes

The results of the seascape genomics analysis were then used to predict the probability of the presence of adaptive genotypes at the scale of the whole Ryukyu Archipelago (Fig. 2B). For each SGEA, the SamBada approach provides parameters of a logistic regression that links the probability of occurrence of the genotype with the value of the environmental variable (Stucki et al., 2017). These logistic models can therefore be used to estimate the probability of presence of the genetic variant for any value of the environmental variable (Joost, 2006; Rochat, E., Leempoel, K., Vajana, E., Colli, L., Ajmone-Marsan, P., Joost, 2016). We assumed that potentially adaptive genotypes could reach any reef of the region since several studies reported migration events between distant islands of the Archipelago (Nakajima et al., 2010; Nishikawa, 2008; Shinzato et al., 2015). Finally, these probabilities where combined into an average probability (i.e. the arithmetical mean) of carrying adaptive genotypes (P_a_) at each reef of the Ryukyu Archipelago.

### Sea current data

Daily records of sea surface current were retrieved from publicly available databases (zonal and meridional surface velocities from the *global-reanalysis-phy-001-030* product; EU Copernicus Marine Service, 2017) and used to compute the direction and speed of currents in the R environment using the *raster* library. By using the resample function of the R *raster* library, we increased resolution of these data from original 0.083° (∼9.2 km) to 0.015° (∼1.6 km) and corrected land pixels (*i*.*e*. removing sea current values) using an high-resolution bathymetry map (Ryan et al., 2009). These day-by-day records of sea currents (from 1993 to 2010) were then stacked to retrieve, for each pixel of the study area, the cumulated speed toward each of the eight neighbouring pixels. For every pixel, the cumulated speed in each of the eight directions was divided by the total speed (the sum of the eight directions) to obtain the probability of transition in each direction (the conductance). This information was used to calculate dispersal costs (the inverse of the square conductance) and was summarized in a transition matrix in the format of the R *gdistance* package (v. 1.2, van Etten, 2018). For the connectivity analysis (Fig. 2C), the transition matrix was used to calculate sea distances (*i*.*e*. the least-cost path) between sampling sites of the genotyped colonies. For the connectivity predictions (Fig. 2D), we calculated the sea distances between all the reefs of the study area, whose coordinates were the centroids of the reef-cells computed as described in the environmental variables section. Importantly, for each pair of reefs (for instance reef_1_ and reef_2_) two sea distances were computed, one for each direction (*i*.*e*. from reef_1_ to reef_2_ and from reef_2_ to reef_1_).

### Connectivity analysis

Genetic distances between sampling sites were calculated using the pairwise F-statistics (Fst; Weir & Cockerham, 1984) as implemented in the R *hierfstat* library (v. 0.04; Goudet, 2005). When there is no dispersal constraint between two sub-populations, the related Fst is equal to zero. Conversely, when dispersal is constrained, Fst increases up to a maximum value of one (isolated sub-populations). To avoid bias due to low sample sizes, we only considered sample sites with more than 10 samples each (7 out of 12).

Next, we built a linear model (hereafter referred to as the connectivity model) to estimate Fst from the sea distances between each pair of sample sites. Since each pair of sites was connected by two distances (from reef_1_ to reef_2_ and *vice versa*), we built three models based on shortest, longest and average sea distance between each pair. As an additional comparison, we also built a connectivity model using Euclidean distances of coordinates as independent variable while maintaining Fst as response variable. For each model, we then ran a t-test to check for the statistical significance of the relationship between genetic and geographical distance. Next, we compared the four models by relying on the coefficients of determination (R2) and the Akaike information criterion (AIC; Bozdogan, 1987) to identify the best connectivity model to use in connectivity predictions.

### Connectivity Predictions

The best connectivity model defined in the connectivity analysis was used to calculate the predicted Fst (pFst) between all pairs of reefs of the Ryukyu Archipelago. To this end, we translated sea distances in pFst using the connectivity model trained during the connectivity analysis. Importantly, since there are two sea distances connecting each pair of reefs, two pFst were computed as well (Fig. S5). Based on these predictions, we were able to calculate two indices for each reef-cell:

- *outbound connectivity index* (OCI; Fig. 4a): the sum of the area (in km^2^) of all the reefs connected under a specific pFst threshold <u>from</u> the focal reef.
- *inbound connectivity index* (ICI; Fig. 4b): the sum of the area (in km^2^) of all the reefs connected under a specific pFst threshold <u>to</u> the focal reef.

These connectivity indices and their interpretation are subordinate to the pFst threshold applied in the calculation. For this reason, it is crucial to set this threshold by considering the size of the study area and the distribution of the pFst values observed (Fig. S5). In this work, we set the pFst threshold to 0.02. In fact, a smaller pFst (for ex. 0.01; Fig. S5) would have informed on local connectivity only (within neighbouring islands) and neglect connectivity at the scale of the Ryukyu Archipelago. In contrast, a higher pFst (for ex. 0.05, Fig. S5) would have exceeded the study area boundaries, causing bias (border effects) in the calculation of the indices for reefs of the southern Islands (Yaeyama and Miyako) of the Archipelago.

### Evaluation of the adaptive potential

The adaptive potential was evaluated by combining the predictions of the presence of adaptive genotypes (P_a_) and connectivity patterns (ICI) in an *index of adaptive potential* (API, Fig. 2E). Indeed, API is a special case of ICI calculated as the sum of the weighted area (in km^2^) of all the reefs connected under a specific pFst threshold to the focal reef. The weight applied to each reef corresponded to the probability of carrying adaptive gentoypes (P_a_). For the pFst threshold, we used the same value (0.02) as employed in the ICI and OCI calculations.

## Results

### Seascape Genomics

We detected 6 significant genotype-environment associations (SGEA, q_G_ and q_W_<0.001, Tab.1) spanning across 6 distinct scaffolds of the *A. digitifera* reference genome. Among them, one SGEA was related to SST variation in April and the others to degree heat week (DHW) frequency.

**Table 1.**
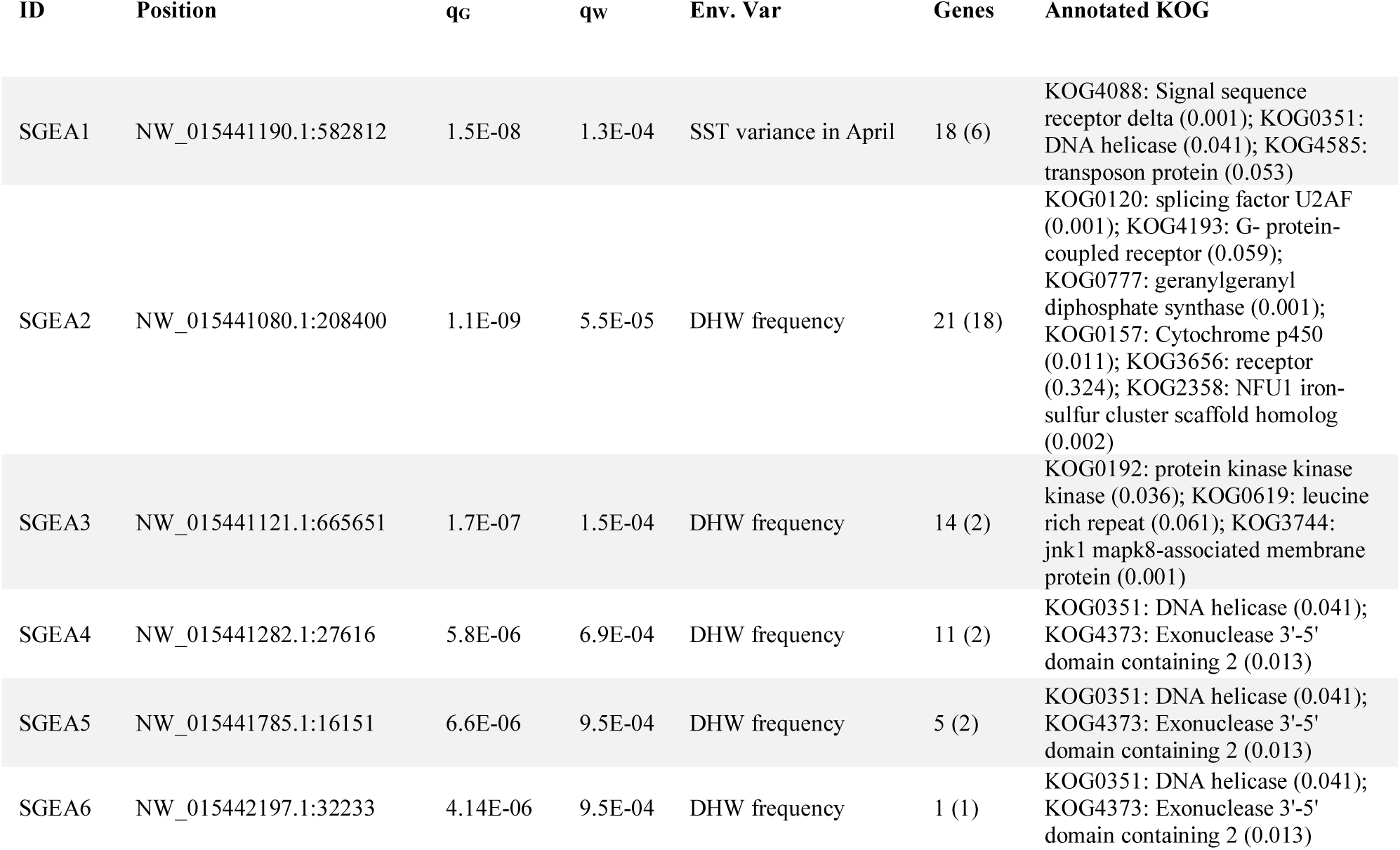
Significant genotype-environment associations. The seascape genomics analysis using the SamBada method detected 6 significant (qG and qW<0.001) genotype-environment associations (SGEA). This table shows, for each SGEA, the genomic position of the concerned SNP (in the format scaffoldID:position; Position), the q-values related to the G-score (qG) and the Wald-score (qW) of the association model, the concerned environmental variables (Env. Var.), the number of predicted genes in a ± 250 kb window around the concerned SNP (with the number of predicted genes annotated by similarity with a known gene, in brackets) and the related annotations on the eukaryote cluster of orthologous genes (KOG; with the frequency of the annotation across the reference genome, in brackets).

SGEA1 described the relationship between SST standard deviation in April and one SNP located within a genomic window (± 250 kb) counting 18 predicted genes, 3 of which annotated to the eukaryotic cluster of orthologs (KOGs) *Signal sequence receptor delta, DNA helicase, transposon protein* (Tab. 1, Box S1). All the remaining SGEAs (SGEA2-6) were related to DHW frequency only. In SGEA2, the concerned SNP was surrounded by 21 predicted genes, 6 of which annotated with KOGs *splicing factor U2AF, G-protein-coupled receptor, geranylgeranyl diphosphate synthase, Cytochrome p450, receptor, NFU1 iron-sulfur cluster scaffold homolog* (Tab.1, Box S2). In SGEA3, the SNP was surrounded by 14 predicted genes, with 3 KOGs annotated as *protein kinase, leucine rich repeat* and *jnk1 mapk8-associated membrane protein* (Tab. 1, Box S3). In SGEA4, 5 and 6, the SNPs were located in genomic windows counting 11, 5 and 1 predicted genes, respectively, and each containing the same two KOGs: *DNA helicase* and *Exonuclease 3’-5’ domain containing 2* (Tab. 1; Box S4, S5, S6).

### Probability of Presence of Adaptive Genotypes

The SGEAs of the seascape genomics analysis were then used as the starting point for predicting the probability of presence of adaptive genotypes (P_a_) across the reefs of the region. Since most of the putative adaptive genotypes were found to be associated with DHW frequency (SGEA2-6; Tab. 1), we chose this variable as the environmental pressure of interest. The average of P_a_ ranged from 0 to 1 (Fig. 3). In Ryukyu Archipelago, P_a_ was higher in Miyako 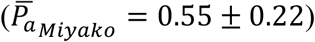 and Okinawa 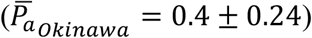, lower in Amami 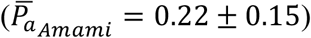 and Yaeyama 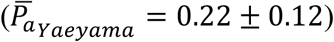, and close to zero in the north of the region (Tokara and Osumi; 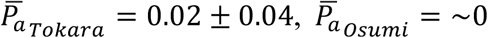; Fig. 3) Outside the Ryukyu Archipelago, a high P_a_ (>0.7) was predicted in northern Philippines while reefs around Taiwan displayed in general low P_a_ (<0.4; Fig. 3).

**Figure 3.**
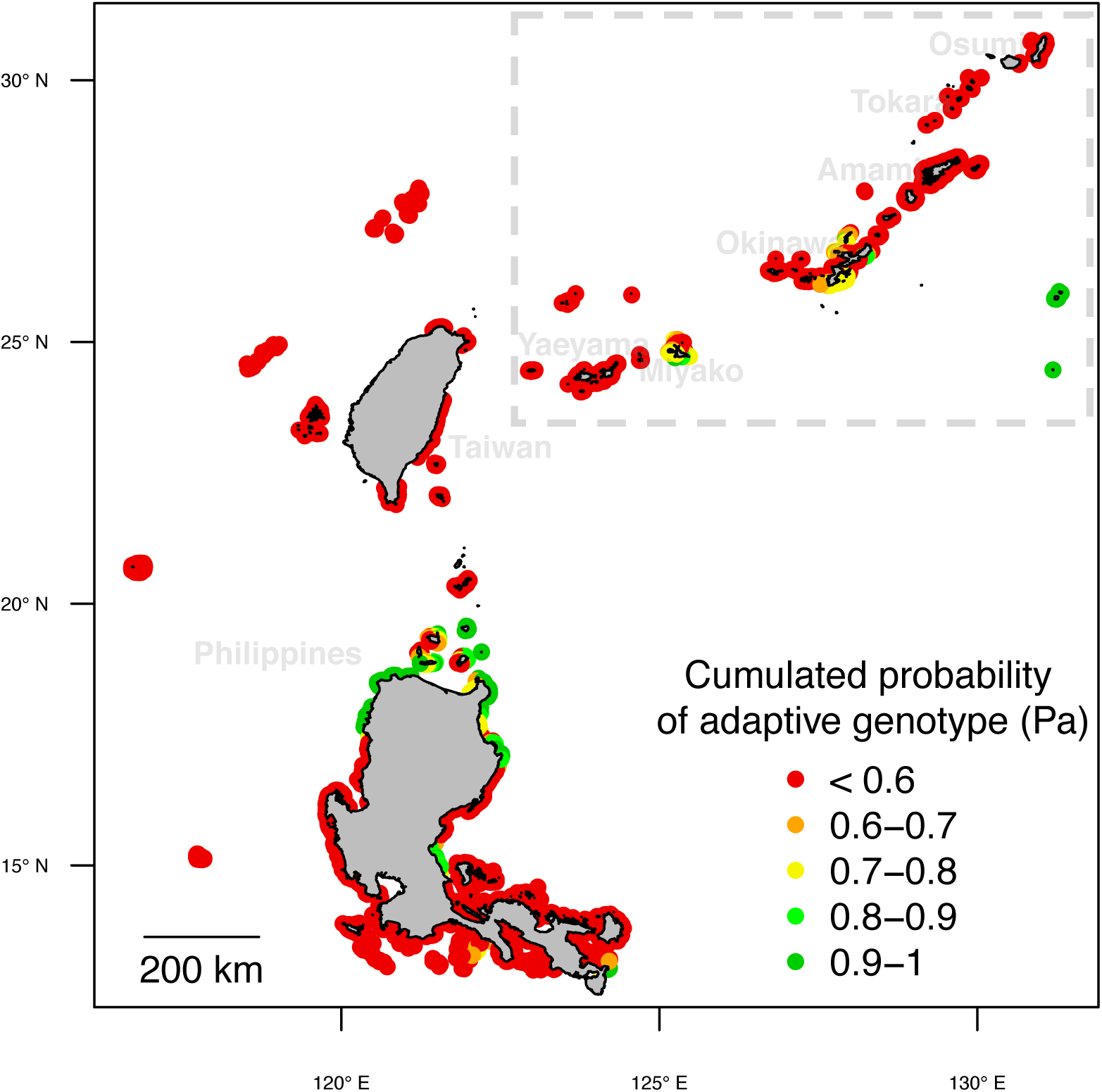
Cumulated probability of carrying adaptive genotypes vs DHW. The map shows the probability of presence of genotypes expected to be linked to adaptation to thermal stress across the study area and the neighbouring regions. Five gene-environment associations (SGEA2-6, Tab. 1) describing the association between distinct putative adaptive genotypes and DHW frequency were used to predict expected genotype frequencies. These expected frequencies were then averaged to compute the cumulated probability of adaptive genotypes. The dashed box highlights the position of the Ryukyu Archipelago.

**Figure 4.**
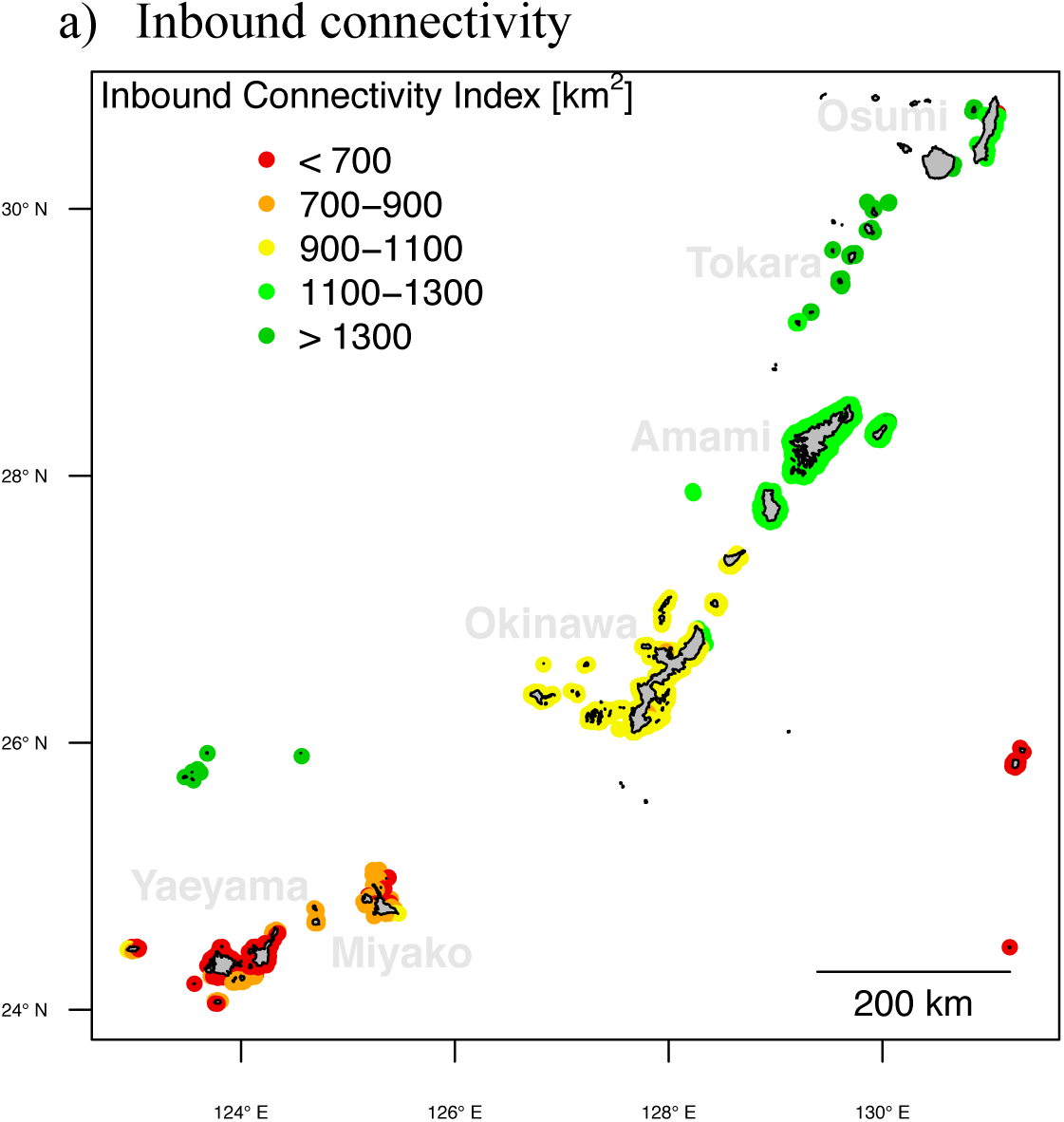
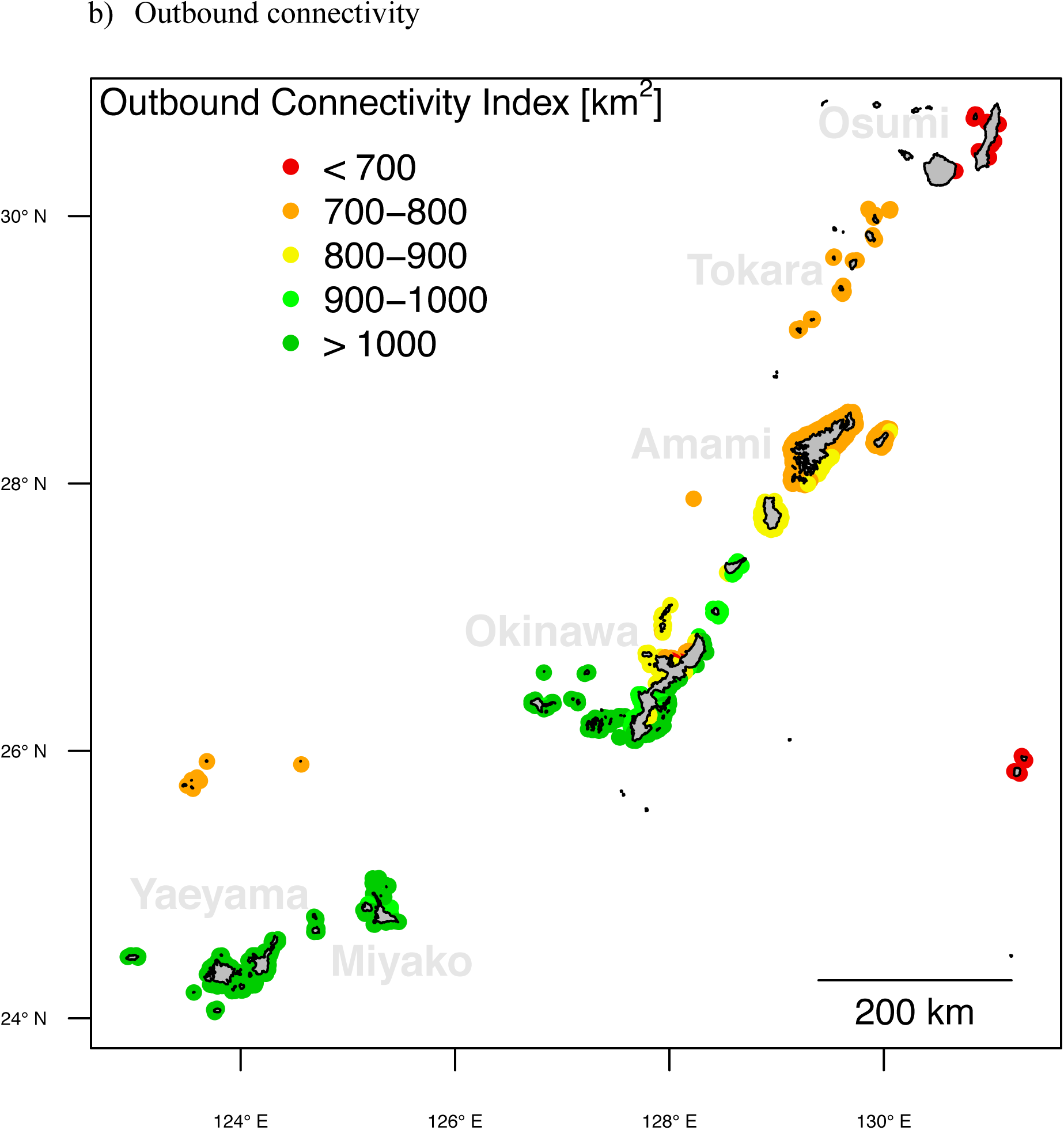
Connectivity indices. The maps show the potential connectivity to (a) and from (b) every reef of the Ryukyu Archipelago. In (a), the inbound connectivity index (ICI) represents the total area (in km^2^) of the reefs that are connected to the focal reef with a pFst<0.02 (pFst toward the focal reef). In (b), the outbound connectivity index (OCI) displays the total area of the reefs that are connected from the focal reef with a pFst<0.02 (pFst from the focal reef).

### Connectivity modelling

The connectivity model based on the shortest sea distances between pairs of sites was the best to describe Fst variation (R2=0.72, AIC=-234; Fig. S4a), compared to models based on the longest sea distance (R2=0.62, AIC=-227, Fig. S4b), the average sea distance (R2=0.66, AIC=- 230, Fig. S4c), and the aerial distance (R2=0.66, AIC=-230, Fig S3d). The connectivity indices were therefore computed using the connectivity model based on the shortest sea distance between sites.

The ICI variation followed a north to south decrease (Fig. 4a). The reefs around the islands in the north of the archipelago (Osumi, Tokara and Amami) were generally those with the highest ICI 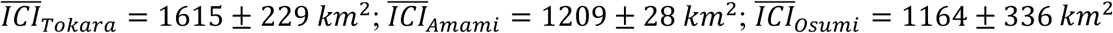; Fig. 4a). In the central area (Okinawa), ICI was lower 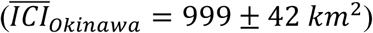, while the lowest ICI values were observed in the southern area (Yaeyama and Miyako; 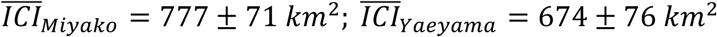; Fig. 4a).

With regards to OCI, we observed a decrease in index with latitude (Fig. 4b). OCI was highest in the southern half of the archipelago (Yaeyama, Miyako and Okinawa; 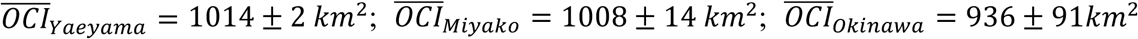; Fig. 4b). A lower OCI was observed in the northern part (Amami and Tokara; 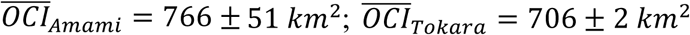; Fig. 4b), while the extreme north of the Archipelago (Osumi) had a very low OCI (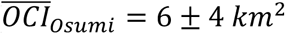; Fig. 4b).

### Evaluation of the adaptive potential

The variations of API were generally structured along the latitudinal axis (Fig. 5). Reefs in the northern part of the Archipelago (Tokara, Amami and Osumi) generally showed the highest API values (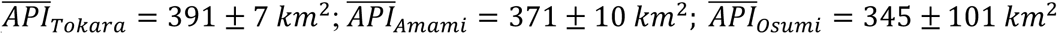; Fig. 5). In the central part of the study area (Okinawa), API was lower 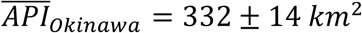; Fig. 5) and in the southern part the lowest API values were observed, (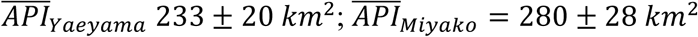; Fig. 5).

**Figure 5.**
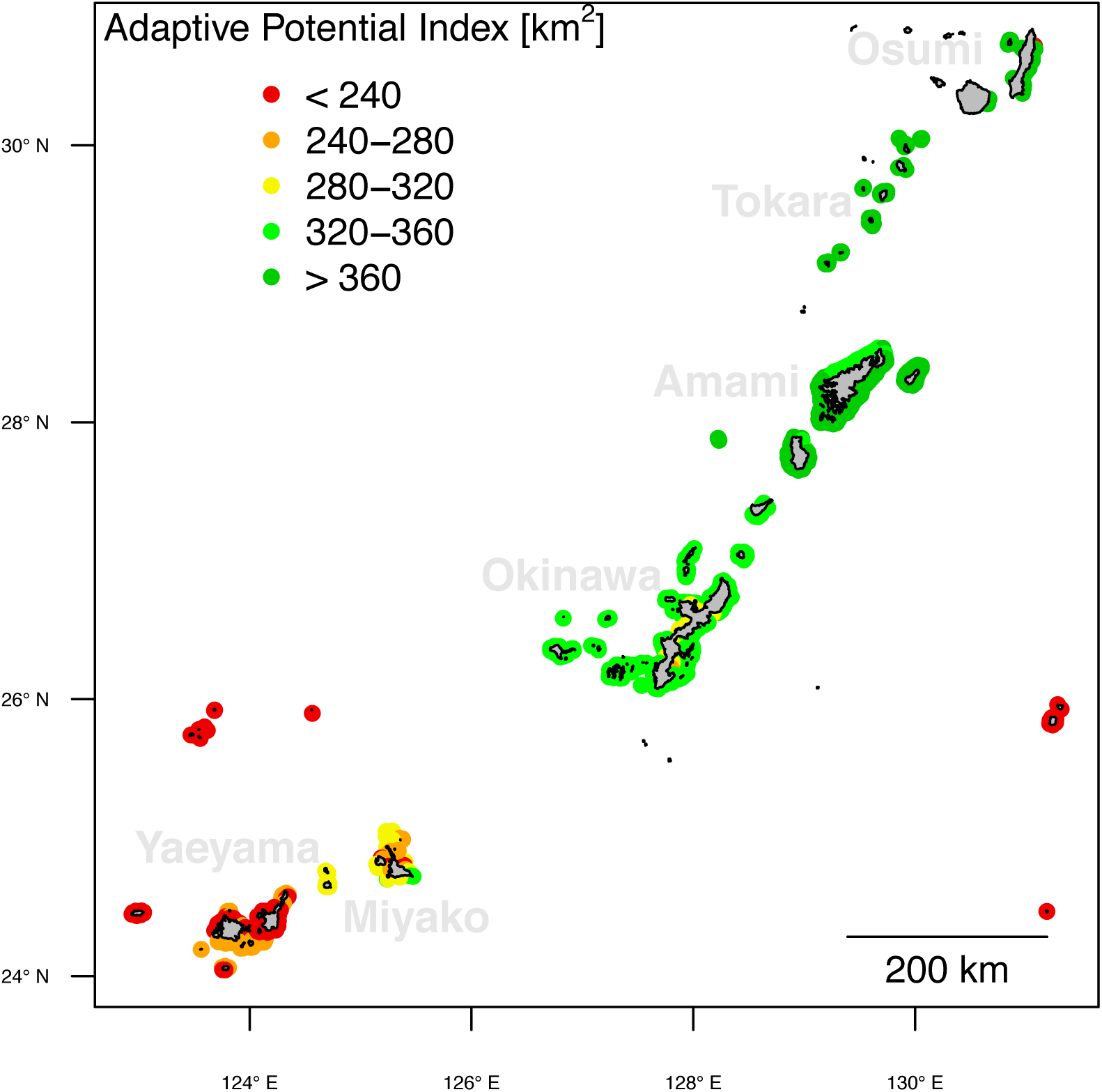
Index of adaptive potential. The map displays the index of adaptive potential (API) against thermal stress (DHW frequency) for every reef of the study area. This index represents the sum of weighted areas of reefs connected to the focal reef with a pFst<0.02 (pFst toward the focal reef). The weight applied correspond to the probability of presence of adaptive genotypes (Pa).

## Discussion

### Adaptation to thermal stress

Thermal stress is expected to be one of the major threats to coral reef survival, where the research for adaptive traits is becoming of paramount importance (Baums, 2008; Logan et al., 2014; Maina et al., 2011). In the present study, the seascape genomics analysis of *A. digitifera* of the Ryukyu Archipelago revealed the presence of 6 genomic regions hosting genetic variants that might confer a selective advantage against thermal stress (Tab. 1). None of the SNPs related to the SGEA lay directly within a coding sequence of a putative gene, but this is rarely the case for causative-mutations (Brodie et al., 2016). In fact, genetic variants in intergenic regions that play a modulatory action on the expression of neighbouring genes are more frequent and can influence loci at a distance of 1-2 Mb (Visel et al., 2009). The fragmentation of the reference genome forced us to limit our search window to ±250 Kb around each SNP, yet we still found annotations corroborating a response to heat stress.

The SNP in SGEA3 was found to be related to KOG3744 (*jnk1 mapk8-associated membrane protein;* Tab. 1). This KOG is rare across the genome of *A. digitifera* (with an expected frequency of 0.0009 per 500 kbs window) and previous research corroborates the hypothesis that this gene plays a role in thermal adaptation. Mitogen-activated-protein kinases (MAPKs) are proteins known to be involved in cellular responses to stress across a range of taxa (Neupane et al., 2013), and the c-Jun-N-terminal kinase (JNK) has previously been shown to be activated under thermal stress in the coral *Stylopohora pistillata* (Courtial et al., 2017).

In SGEA1, 4, 5 and 6, one KOG recurred in the annotations: KOG0351 (*DNA helicase*; Tab. 1). The expected frequency of this KOG is 0.04 per 500 kbs window and remarkably we found 4 of them in 6 distinct 500 kbs windows around SGEA associated with thermal stress. These orthologs annotate a particular type of DNA helicases (swissprot IDs: Q91920, Q14191) known as “helicases Q” or “RecQ” (Box S1, 5-7), which are involved in the DNA repairing mechanism caused by UV-light damage in prokaryotes (Courcelle & Hanawalt, 1999), and for which light-stress driven effects were observed in eukaryotic cells as well (Sharma et al., 2006). The modulation of this mechanism might therefore play a role in increasing *A. digitifera* resistance against light-stress associated with heat waves.

Seascape/Landscape genomics studies are susceptible to high false discovery rates, especially when neutral genetic structure is not accounted for (Rellstab et al., 2015). To take this element into account, we processed data following a conservative pipeline and used models explicitly integrating demographic processes. We also set a restrictive threshold to filter significant association-models (*i*.*e*. q<0.001 in two distinct tests). Most of the SGEA found were linked to DHW, most probably because this variable represents one of the main constraints to coral survival (Hughes et al., 2003). It is also possible that the initial application of a sampling scheme adapted to seascape genomics (unlike the one used by Shinzato et al. 2015 who did not consider environmental variability) would have increased sensitivity to other types of adaptive signals (Riginos et al., 2016; Selmoni et al., 2019).

We found SGEA related to thermal stress displaying annotations corroborating a role in cellular heat responses. Nonetheless, annotations analysis is only a first step in the validation of the SGEA detected. Ideally, further assays as reciprocal transplantations (Palumbi et al., 2014), experimental conditioning (Krueger et al., 2017) and molecular analysis (Courtial et al., 2017) could ascertain and help quantifying the link between each genotype and the putative selective advantage it might confer.

### Connectivity patterns

Coral dispersal is driven by the water flow (Paris-Limouzy, 2011), which is highly asymmetrical in this region (north-east oriented) due to the Kuroshio current (Nishikawa, 2008). As previously observed, the main patterns of migrations in this population occurs from the south-west to the north-east (Shinzato et al., 2015). Reefs in the southern part of the study area (Yaeyama and Miyako) showed the lowest ICI values (Fig. 4a), suggesting a potential lack of recruits arriving from other reefs of the region. In fact, the genetic diversity of southern reefs of the Ryukyu Archipelago is likely to depend on the recruits arriving from the east-coast of Taiwan and the northern Philippines, which are located upstream of the Kuroshio current (Fig. S5a; Chen & Shashank, 2009).

In the previous study on this data (Shinzato et al., 2015), reefs from Yaeyama resulted as those with the lowest heterozygosity rates across the study area. This observation was attributed to a population bottleneck caused by the 1998 bleaching event, but it is worth noting that reefs on the west coast of Okinawa showed higher heterozygosity rates despite having suffered recurrent bleaching events since 1998 (Donner et al., 2017). The lower heterozygosity rates in Yaeyama therefore might reflect not only the effects of past bleaching, but also the relative isolation of these islands from the reefs of the region (Fig. 4a).

In line with the same previous observations (Shinzato et al., 2015), the OCI value showed (Fig. 4b) that the southern reefs (Yaeyama and Miyako) are those expected to be the most prominent source of recruits for the rest of the Archipelago. Given this crucial aspect, it is even more important to preserve southern reefs of the Ryukyu Archipelago from the risks of isolation (e.g. inbreeding depression; Keller & Waller, 2002).

### Adaptive potential in the 2016 bleaching event

Reefs from islands of Miyako, Okinawa and the northern part of Amami were those most likely to carry adaptive genotypes against thermal stress (Fig. 3). Previous work reported severe bleaching in Okinawa in 1998 (Yamazato, 1999) and that adapted colonies might have resisted (Van Woesik et al., 2004). In contrast, reefs in the northern part of the Archipelago (Amami, Tokara and Osumi) experienced bleaching with moderate severity during the 1998 event (Donner et al., 2017), which might explain why adaptive genotypes are not expected at the same frequency (Fig. 3).

The adaptive potential index (API) defines the convergence between the probability of carrying adaptive genotypes with connectivity predictions (Fig. 5). Reefs in the northern part of the Archipelago (Amami, Tokara and Osumi) showed a higher API compared to those in the southern half of the region (Okinawa, Yaeyama and Miyako). Two reasons may explain this result: 1) these northern reefs are located downstream (on the Kuroshio Current) of two areas where putative adapted reefs are frequent (Okinawa and Miyako; Fig. 3); 2) the region of Northern Philippines, hosting high density of putative adapted reefs (Fig. 3), is more connected to the northern part of the Ryukyu Archipelago than with the southern part (Fig. S6). This may also explain why, despite hosting putative adapted reefs (Fig, 3), the Miyako area showed among the lowest API values of the Archipelago (Fig. 5).

In 2016, the first mass bleaching event occurred in Japan since the Shinzato and colleagues published the genetic data re-analysed in this work (Kimura et al., 2018). Field surveys related to this bleaching event reported severe bleaching in Yaeyama (intensity up to 99%, mortality up to 68%) and in Miyako (intensity up to 70%, mortality up to 67%; Tab. 2). In Okinawa and Amami, the impact of this same bleaching event was moderate to mild (Okinawa: intensity up to 48%, mortality up to 13%; Amami: intensity 8% and mortality 2%; Tab. 2). Reefs predicted to show low API (the southern reefs) appeared to suffer more severe bleaching than those in the northern region (which showed higher API; Fig. 5), but care must be taken in the interpretation due to the confounding role of the degree of heat stress (Tab. 2).

**Table 2.** Field report of the 2016 mass bleaching event. The table shows the severity and mortality associated with the 2016 bleaching event as reported by Global Coral Reef Monitoring Network (Kimura et al., 2018). For every region surveyed in this report (identified by an ID and a region name), we show the corresponding region in our study and the associated average API, and degree of heat stress in 2016 (estimated as the number of days with DHW>4°C). Colours scales highlight the value variations of each column.

| ID | Region Name | Region Area<br>(this study) | Bleaching [%] | Mortality [%] | API [km<sup>2</sup>] | $P_a$ | Heat Stress<br>[# of days] |
| --- | --- | --- | --- | --- | --- | --- | --- |
| 3 | Amami Islands | Amami | 8.5 | 2.1 | 371 | 0.26 | 68 |
| 4 | Okinawa Island, East coast | Okinawa | 21 | 0.7 | 339 | 0.60 | 69 |
| 5 | Okinawa Island, West coast | Okinawa | 13.1 | 4.3 | 328 | 0.36 | 75 |
| 6 | Okinawa Outer Islands | Okinawa | 48.4 | 13.5 | 336 | 0.70 | 82 |
| 7 | Kerama Islands | Okinawa | 7.3 | 5.4 | 334 | 0.08 | 81 |
| 9 | Miyako Island | Miyako | 68.8 | 31 | 282 | 0.61 | 84 |
| 10 | Miyako Outer Reefs | Miyako | 70.1 | 67.5 | 293 | 0.69 | 85 |
| 11 | Ishigaki Island, East coast | Yaeyama | 47.9 | 8.8 | 230 | 0.38 | 81 |
| 12 | Ishigaki Island, West coast | Yaeyama | 63.2 | 14.8 | 226 | 0.11 | 80 |
| 13 | Sekisei Lagoon, North | Yaeyama | 91.5 | 46.9 | 225 | 0.16 | 81 |
| 14 | Sekisei Lagoon, East | Yaeyama | 99.5 | 67.9 | 238 | 0.29 | 82 |
| 15 | Sekisei Lagoon, Center | Yaeyama | 94.9 | 49.7 | 241 | 0.24 | 82 |
| 16 | Sekisei Lagoon, South | Yaeyama | 98.2 | 50 | 254 | 0.19 | 83 |
| 17 | Iriomote Islands | Yaeyama | 94.3 | 34.8 | 236 | 0.28 | 80 |

Indeed, satellite records of sea temperature (EU Copernicus Marine Service, 2017) show that in 2016 the number of days in temperature anomaly (DHW>4°C; Liu et al., 2003) was higher in the southern part of the Archipelago (Yaeyama: ∼81 days; Miyako: ∼84 days) than in the northern region (Okinawa: ∼76 days; Amami: ∼68; Tab. 2). Nevertheless, when two distant sites had a comparable degree of heat stress, higher API was generally associated with a reduced severity in bleaching. For instance, reefs in Kerama (Okinawa) and Sekisei Lagoon North (Yaeyama) both suffered 81 days of heat stress in 2016, but the bleaching intensity in the Sekisei Lagoon was more than ten times higher than observed for Kerama (91% vs 7%), with a lower API (225 km^2^ vs 334 km^2^; Tab 2).

While these field observations seem to corroborate our predictions on adaptive potential, it is important to consider that they do not refer specifically to *A. digitifera*, but to the coral community as a whole (Kimura et al., 2018). Additionally, other local stressors (for instance anthropogenic pollution) might have modulated the bleaching response (Ateweberhan et al., 2013). Future bleaching surveys, with larger sample sizes and bleaching data referring to the specific coral genus, might provide a more reliable ground for validating our predictions.

### Application in conservation

Conservation policies require objective and quantifiable information to prioritize areas for intervention efforts (OECD, 2017). In this study, we presented an original framework to calculate indices matching these requirements to describe the connectivity and adaptive potential against thermal stress of a flagship coral species of the north-western Pacific. Insights of this kind are essential for effective planning of coral conservation strategies (Baums, 2008; Logan et al., 2014; Palumbi, 2003; van Oppen et al., 2015b).

As they are derived from a universal metric of population connectivity (Fst; Weir & Cockerham, 1984), the indices we propose here are computable for any coral species. Thus, connectivity indices for different species can be compared or aggregated for conservation management planning within a region. Furthermore, each of the indices we propose is expressed in a tangible spatial unit (km^2^) that allows for comparison between different datasets and areas.

As an example, the predictions from the connectivity indices can be used to support the planning of marine protected areas (MPAs; Box 1a). An ideal placement of an MPA should ensure that the connectivity to the rest of the reef system is optimal (Krueck et al., 2017; C. J. Thomas et al., 2014), and the OCI provides this information (Fig. 4b). Furthermore, the computation of the ICI (Fig. 4a) from a protected area to the rest of the reef system could be used to compare how different locations of MPAs may modify the connectivity to other specific regions (Box 1a).

#### Box 1.

Examples of applications of ICI (a) and API (b) in conservation management

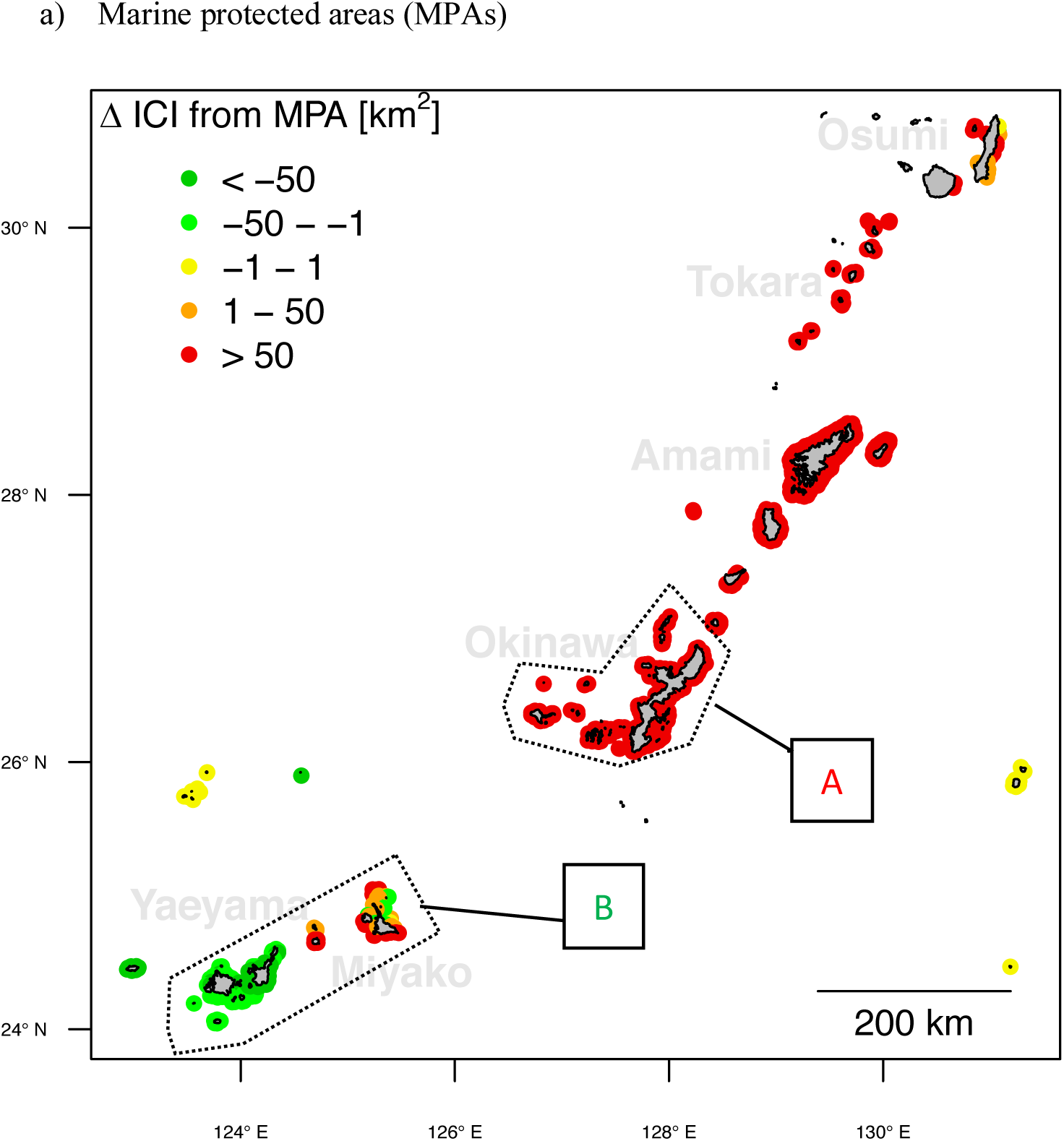

Two different locations for MPA of the same size are compared. In the first location (A), the MPA is established in Okinawa, and in Miyako and Yaeyama for the second one (B). For every reef of the Ryukyu Archipelago, we computed the ICI from each of the two MPA locations. In this map, we show the difference of the two ICI (Δ *ICI* = *ICI*_*A*_ – *ICI*_*B*_). An overall 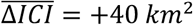 indicates that the MPA in (A) is expected to produce a stronger increase of the inbound connectivity of reefs of the study area. However, these positive effects do not extend to southern reefs (Yaeyama and Miyako), where an MPA in (B) would provide a stronger increase in ICI.

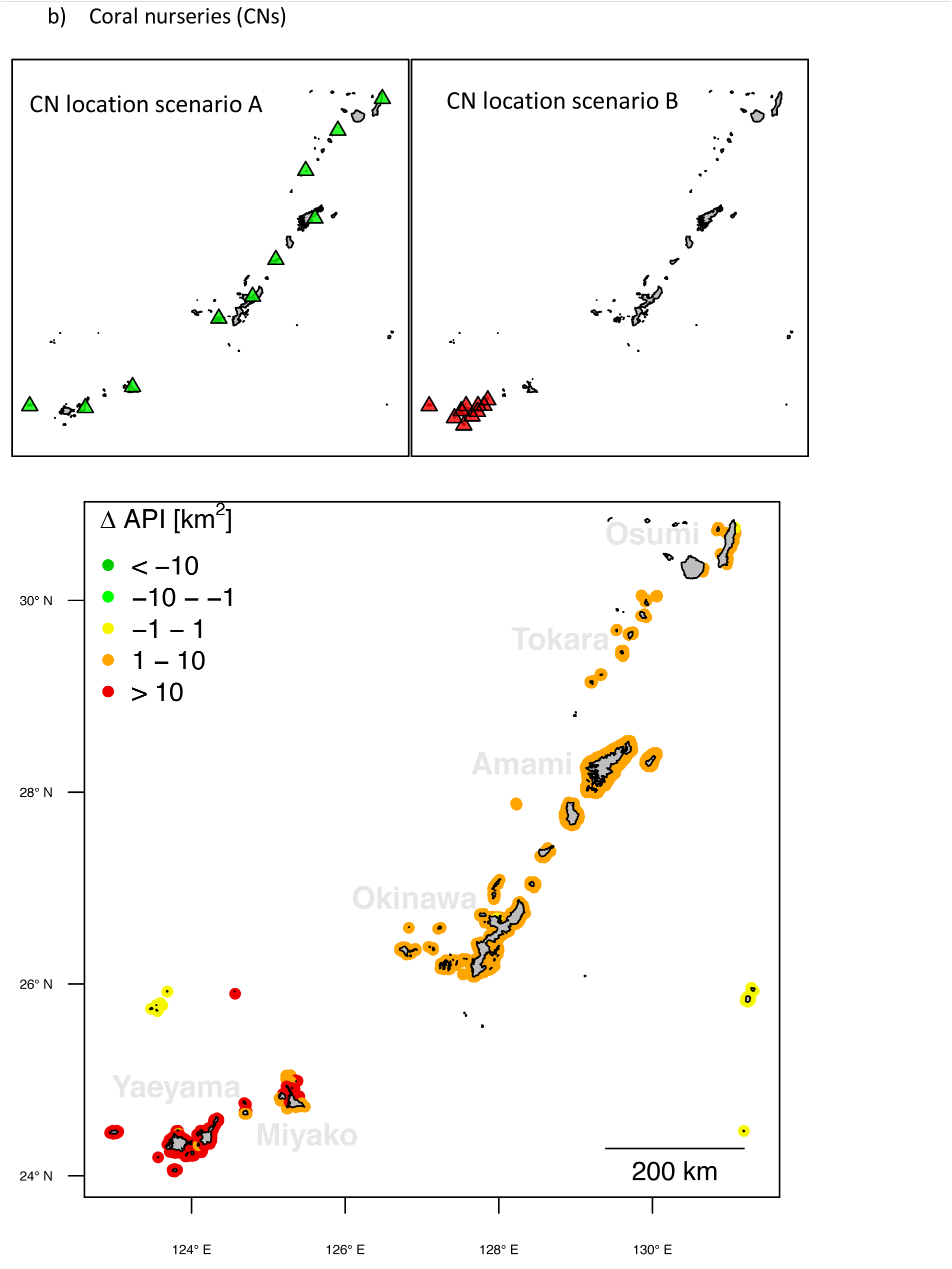

The different locations of two CNs network are compared. Each network is composed of 10 reefs where the P_a_ was set equal to 1 (simulating the restoration via transplantation of adapted colonies). In the first scenario (A, green triangles), CNs are established in the island of Yaeyama. In the second scenario (B, red triangles), CNs are located all across the archipelago. For every reef of the study area, API is calculated under the two network scenarios (A and B). The map shows the difference of API between the two locations (Δ *API*_*A*_ – *API*_*B*_). Overall 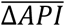 is +5 *km*^2^, therefore suggesting that the CN network A provides a general increase of adaptive potential in the study area. Such positive effects are visible in any region of the Archipelago.

The indices proposed in this work could also support coral nursery establishment plans (Box 1b). In this type of conservation strategy, the transplantation of adapted colonies (*i*.*e*. those carrying the adaptive genotypes) can be used to reinforce the adaptive potential of a population (Baums, 2008; van Oppen et al., 2015b). Conservation managers can simulate the transplanting procedure by setting P_a_=1 at reefs where the nurseries will be established and then measure how these changes would impact the API of the population (Box. 1b).

To date, the calculation of these indices can be performed using the R scripts and codes (R Core Team, 2016) made publicly available in this research. In the future, however, this framework should be transposed to a more user-friendly interface to facilitate its use by conservation managers.

## Conclusions

This study highlights the value of a seascape genomics approach for supporting the conservation of corals. We applied it to a flagship coral species of the Ryukyu Archipelago and identified genetic variants that may underpin adaptation to thermal stress. Coupling this information with a genetic analysis of connectivity made it possible to evaluate the adaptive potential at the scale of the entire study area. The outputs of this analysis are quantitative indices that are could be used to support objective prioritization of reefs in conservation plans. This framework is transferable to any coral species on any seascape and therefore constitutes a useful conservation tool to evaluate the genomic adaptive potential of coral reefs worldwide.

## Data Archiving Statement

All the data and codes used in this article will be made publicly available on Dryad after manuscript is accepted for publication.

## Acknowledgements

We thank Annie Guillaume and François Bonhomme for the useful comments and suggestions provided during the redaction of this paper.

