## supplementary material for "Seascape genomics as a new tool to empower coral reef conservation strategies: an example on north-western Pacific *Acropora digitifera*"


| Table of content | |
|---|---|
| Tab S1 | **Environmental Variables included in the Seascape Genomics analysis** |
| Fig. S1 | **Fst analysis by genomic position.** |
| Fig. S2 | **Discriminant Analysis of Principal Components (daPCA) of the Genotype Matrix.** |
| Fig. S3 | **Diagram of workflow for computation of genotype-environment association models.** |
| Box S1 | **Genotype Environment Association GEA1.** |
| Box S2 | **Genotype Environment Association GEA2.** |
| Box S3 | **Genotype Environment Association GEA3.** |
| Box S4 | **Genotype Environment Association GEA4.** |
| Box S5 | **Genotype Environment Association GEA5.** |
| Box S6 | **Genotype Environment Association GEA6.** |
| Fig. S4 | **Connectivity models.** |
| Fig. S5 | **Example of pFst variation across study area.** |
| Fig. S6 | **pFst from Northern Philippines.** |

**Supplementary Table 1. Environmental Variables included in the Seascape Genomics analysis.** For each geoinformatics product, the table shows the sources (CMEMS= Copernicus Marine Environment Monitoring System; NEO= Nasa Earth Observations; NOAA= National Oceanic and Atmospheric Administration; IEDA= Interdisciplinary Earth Data Alliance; UNEP = United Nations Environment Program) and the corresponding identifier (Product Name). Temporal range and resolution indicate the duration and the frequency of the remote sensing period, respectively. The variables included in the Seascape Genomics analysis are listed in the *Derived Variables* column, with the number in curly brackets indicating the total number. In total, 315 derived variables were computed.

| **Variable Name** | **Product Name** | **Source** | **Temporal Range** | **Temporal Resolution** | **Spatial Resolution** | **Derived Variables** |
|---|---|---|---|---|---|---|
| Sea Surface Temperature | SST_GLO_SST_L4_REP_OBSERVATIONS_010_011 | CMEMS | 1985-2007 | Daily | 0.05 ° | Sea Surface Temperature means and averages (by month and overall), DHW frequency {27} |
| Sea Surface Salinity | global-reanalysis-phy-001-030-monthly-SSS | CMEMS | 1993-2010 | Monthly | 0.083 ° | Means and averages (by month and overall) {26} |
| Cloud Fraction | Cloud Fraction | NEO | 2000-2010 | Monthly | 0.1 ° | Means and averages (by month and overall) {26} |
| Net Solar Radiation | Net Solar Radiation | NEO | 2006-2010 | Monthly | 0.25 ° | Net Solar Radiation means and averages (by month and overall) {26} |
| Chlorophyll Concentration | oc-glo-chl-multi-l4-gsm | CMEMS | 1998-2010 | Monthly | 4 km | Means and averages (by month and overall) {26} |
| Suspended Particulate Matter | oc-glo-opt-multi-l4-spm | CMEMS | 1997-2010 | Monthly | 4 km | Means and averages (by month and overall) {26} |
| Coloured Dissolved Matter | oc-glo-opt-multi-l4-cdm443 | CMEMS | 1997-2010 | Monthly | 4 km | Means and averages (by month and overall) {26} |
| Sea Currents Velocity | global-reanalysis-phy-001-030 | CMEMS | 1993-2010 | Monthly | 0.083 ° | Means and averages (by month and overall) {26} |
| Photosynthetically Available Radiations | erdMH1par0mday | NOAA | 2003-2010 | Monthly | 4 km | Means and averages (by month and overall) {26} |
| Bathymetry | Global Multi-Resolution Topography | IEDA | - | - | down to 100 m | Depth {1} |
| Population Density | SEDAC | Columbia University | 2015 | - | 0.1 ° | Population density on buffer 50 km {1} |
| Sea Surface Salinity (daily) | global-reanalysis-phy-001-030—daily-SSS | CMEMS | 1993-2007 | Daily | 0.083 ° | In combination with SST: means and average (monthly and overall) for pH, dissolved organic carbon, alkalinity. {78} |

**Supplementary Figure 1. Fst analysis by genomic position.** The fixation index (Fst) between each pair of sub-populations (a: Okinawa and Kerama, b: Okinawa and Yaeyama, c: Yaeyama and Kerama) was calculated for every SNP in the filtered genetic dataset (graphs on the first column). Fst values were then averaged by genomic position: either by contig (graphs on the second column), non-overlapping 100 kbs window (third column) or non-overlapping 50 kbs window (fourth column).

a) Fst Okinawa – Kerama

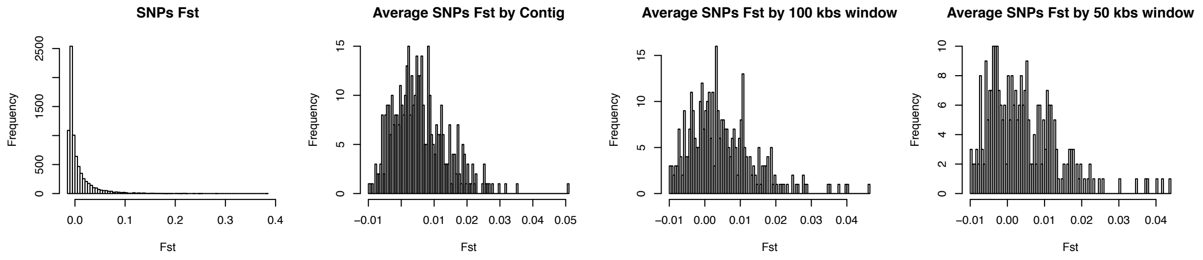


b) Fst Okinawa – Yaeyama

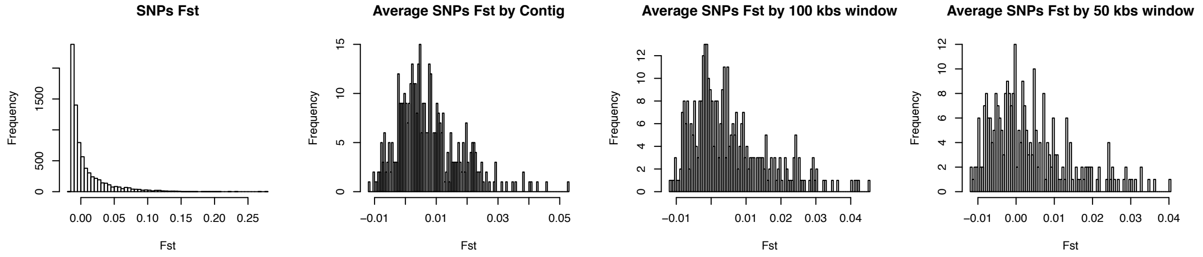


c) Fst Yaeyama – Kerama

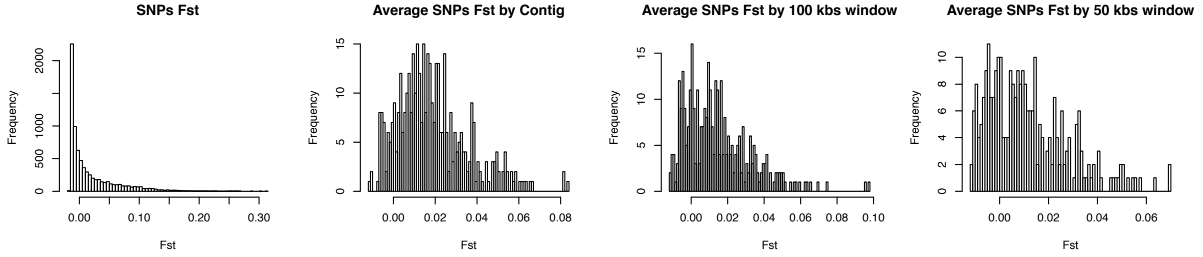

**Supplementary Figure 2. Discriminant Analysis of Principal Components (daPCA) of the Genotype Matrix.** A daPCA of the genotype table was performed to investigate the neutral structure of genetic variation in the population using the R adegenet package. Graph a) shows the (Bayesian Information Criterion) BIC value against the number of clusters, suggesting the presence of two groups. Graph b) displays that the first discriminant function allows to distinguish these groups and the map in c) shows the average of this value across sampling sites. We can see a strong contrast between sites reefs in the south and those in the North of Okinawa islands, together with those in the south of the Archipelago (Yaeyama).

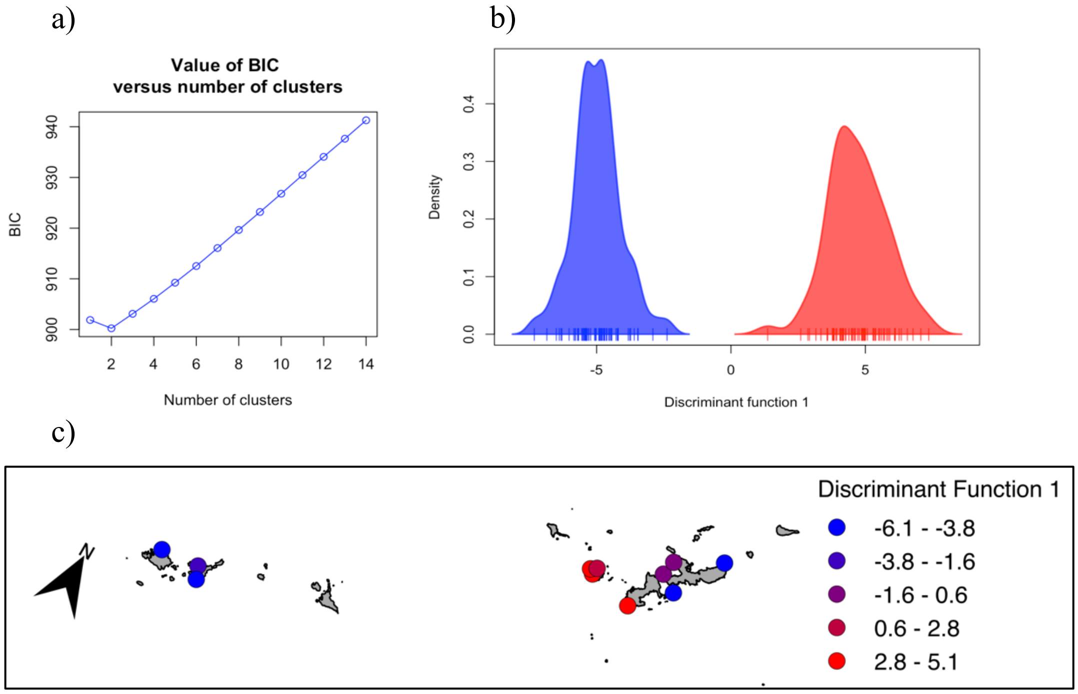

**Supplementary Figure 3. Diagram of workflow for computation of genotype-environment association models.** The environmental variables are summarized in groups of correlated descriptors (A) and one variable per group is randomly chosen for the analysis of association with genotypes. For every SNP in the genotype matrix, the three respective genotypes (C) are investigated for associations with the uncorrelated environmental variables (D). The association models are than ranked by Gscore and deemed as significant if the q-value associated to G-score and Wald-test are below 0.0001 (E). The variables correlated to the one providing the best significant model (in this case SST1, environmental cluster 1) are then used to compute the association-models against all the genotypes of the SNP. The variable providing the best model is identified by ranking the G-scores (in this case DHW). This approach allows to fasten the computation, since all the SNPs are matched with all the environmental clusters, but only the significant ones with all the variables within a cluster.

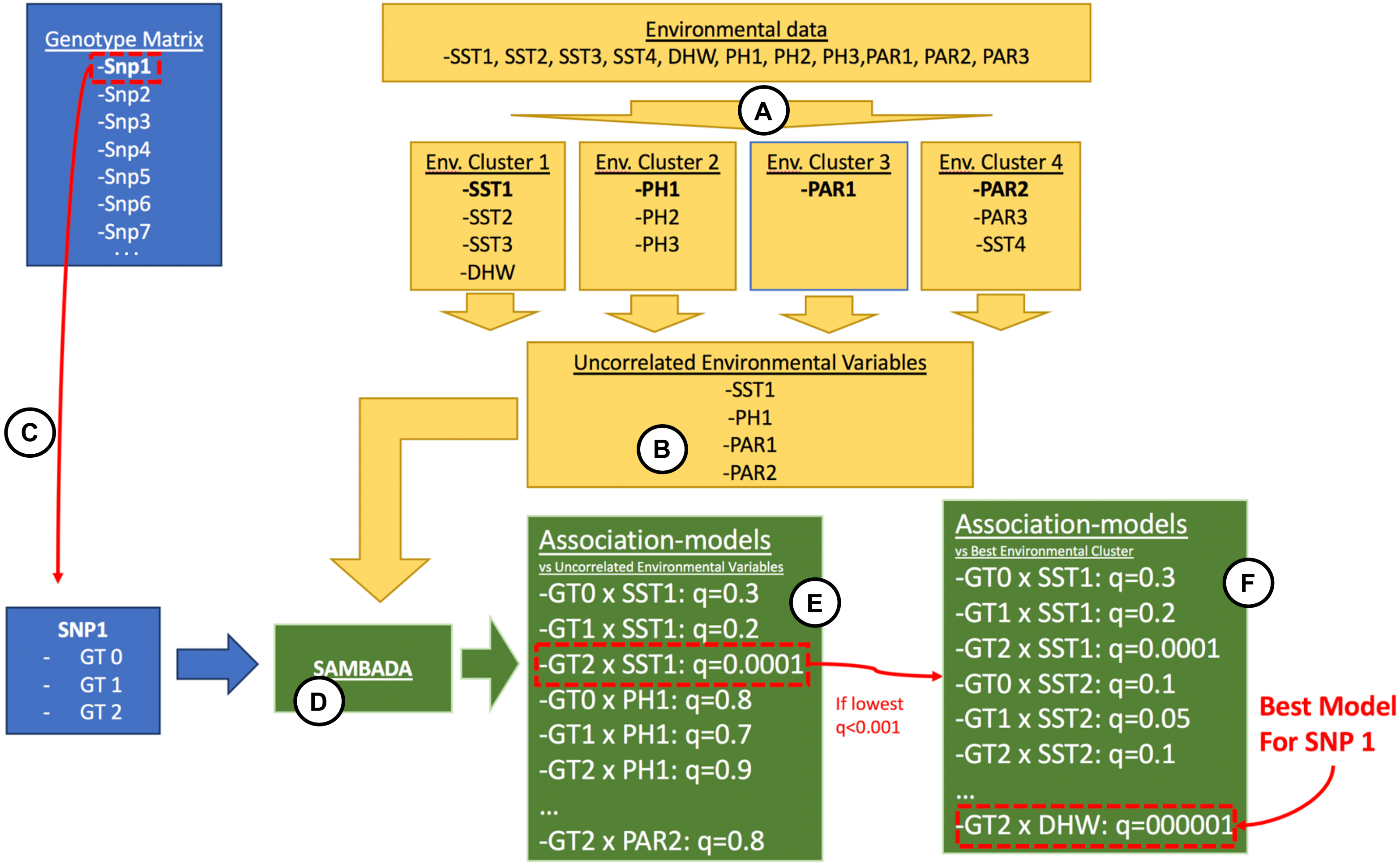

**Supplementary Box 1. Genotype Environment Association GEA1.** Graph in a) shows the logistic model describing the presence (Probability=1) and absence (Probability=0) of the genetic variant as a function of the environmental gradient (SST variation in April). The frequency of the genotype occurrences in real data is represented by the grey bars. The map in b) show the average values of the environmental variable and the genotype frequency by sampling sites. Table in c) illustrates the annotations of the reference genome in the neighborhood (± 250 kb) of the significant SNP. For every locus, the table shows the starting and ending position on the scaffold, the best match of the sequence similarity search against the swissprot database (swissprot, $E<10^{-7}$), the annotation of the best match (annotation) and the related eukaryotic cluster of orthologs (KOGs).

a)

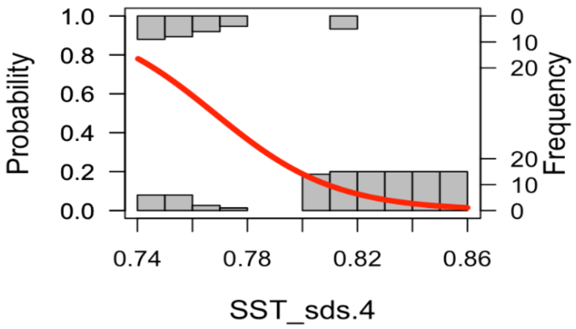


b)

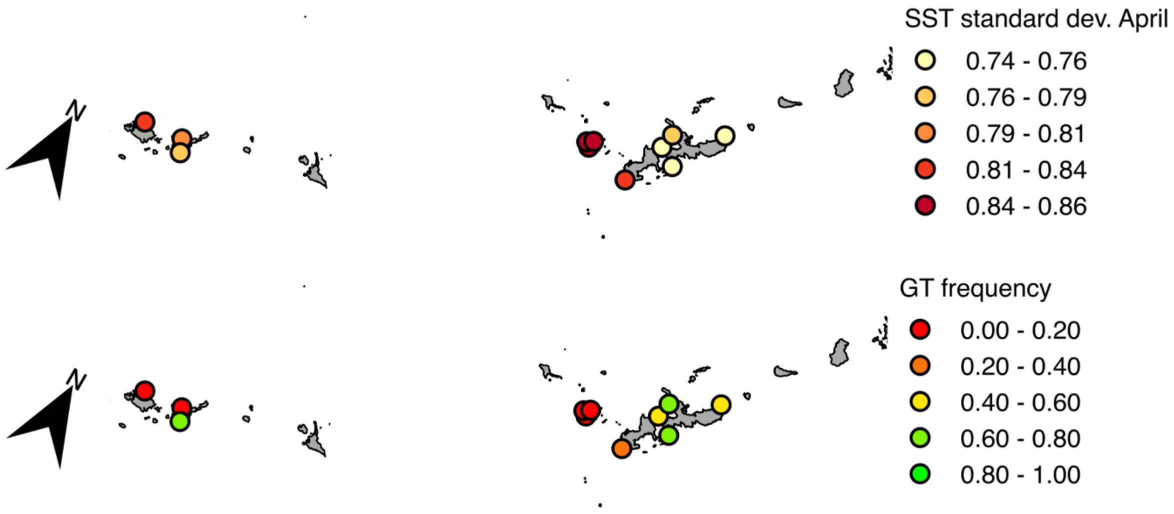


c)

| gene ID | position start | position end | swissprot | annotation | KOGs |
|---|---|---|---|---|---|
| **LOC107339659** | 326653 | 354193 | Q2TBX5 | Translocon-associated protein subunit delta (TRAP-delta) (Signal sequence receptor subunit delta) (SSR-delta) | KOG4088: Signal sequence receptor delta |
| **LOC107339661** | 376771 | 379978 | | | |
| **LOC107339660** | 399170 | 400763 | | | |
| **LOC107339671** | 474958 | 477640 | A4K2W5 | Serine/threonine-protein kinase 4 (EC 2.7.11.1) [Cleaved into: Serine/threonine-protein kinase 4 37kDa subunit (MST1/N); Serine/threonine-protein kinase 4 18kDa subunit (MST1/C)] | |
| **LOC107339672** | 482489 | 485674 | | | |
| **LOC107339625** | 497191 | 498825 | Q96DM1 | PiggyBac transposable element-derived protein 4 | |
| **LOC107339673** | 513832 | 521204 | | | |
| **LOC107339626** | 529212 | 530667 | O93530 | Werner syndrome ATP-dependent helicase homolog (EC 3.6.4.12) (Exonuclease WRN) (EC 3.1.-.-) (Focus-forming activity 1) (FFA-1) | |
| **LOC107339627** | 544539 | 547941 | Q9I920 | Bloom syndrome protein homolog (EC 3.6.4.12) (RecQ helicase homolog) | KOG0351: DNA helicase |
| **LOC107339675** | 556712 | 558640 | | | |
| **LOC107339676** | 558665 | 559637 | Q17QR8 | Putative nuclease HARBI1 (EC 3.1.-.-) (Harbinger transposase-derived nuclease) | KOG4585: transposon protein |
| **LOC107339629** | 578054 | 578395 | | | |
| | | | | **(SNP at 582812)** | |
| **LOC107339677** | 592624 | 608012 | | | |
| **LOC107339630** | 660035 | 660724 | | | |
| **LOC107339631** | 683947 | 684585 | | | |
| **LOC107339632** | 704921 | 706354 | | | |
| **LOC107339678** | 713729 | 718891 | | | |

| | | |
|---|---|---|
| **LOC107339633** | 713796 | 716338 |
| **LOC107339634** | 741458 | 742324 |

**Supplementary Box 2. Genotype Environment Association GEA2.** Graph in a) shows the logistic model describing the presence (Probability=1) and absence (Probability=0) of the genetic variant as a function of the environmental gradient (DHW frequency). The frequency of the genotype occurrences in real data is represented by the grey bars. The map in b) show the average values of the environmental variable and the genotype frequency by sampling sites. Table in c) illustrates the annotations of the reference genome in the neighborhood (± 250 kb) of the significant SNP. For every locus, the table shows the starting and ending position on the scaffold, the best match of the sequence similarity search against the swissprot database (swissprot, E<$10^{-7}$), the annotation of the best match (annotation) and the related eukaryotic cluster of orthologs (KOGs).

a)

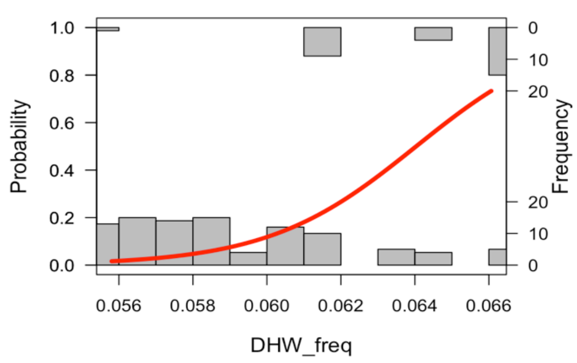


b)

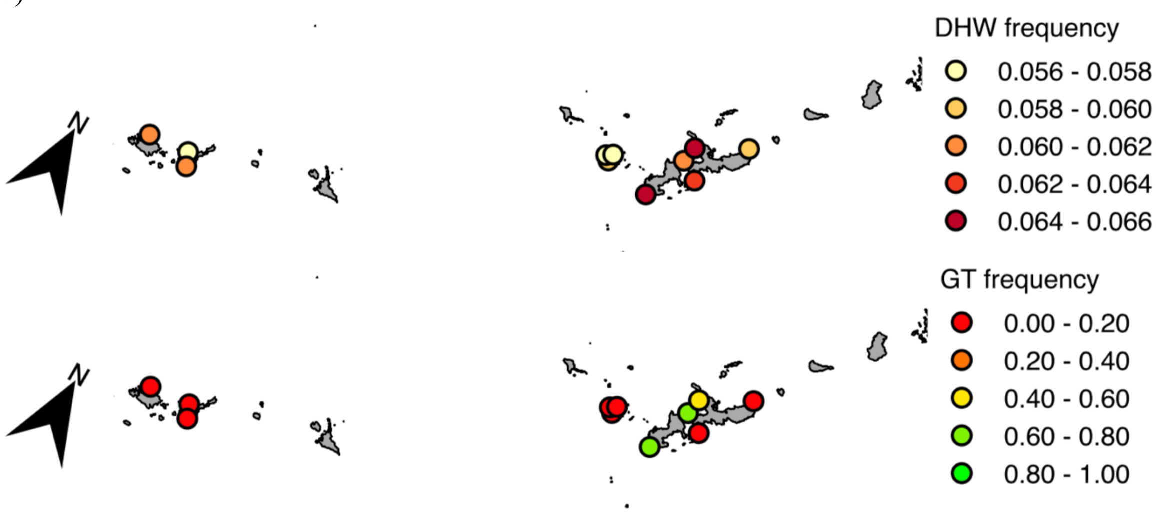


c)

| gene ID | position start | position end | swissprot | annotation | KOGs |
|---|---|---|---|---|---|
| **LOC107329834** | 392 | 11511 | Q24562 | Splicing factor U2AF 50 kDa subunit (U2 auxiliary factor 50 kDa subunit) (U2 snRNP auxiliary factor large subunit) | KOG0120:splicing factor U2AF |
| **LOC107330293** | 26573 | 37385 | Q01528 | Hemagglutinin/amebocyte aggregation factor (18K-LAF) | |
| **LOC107330170** | 34492 | 79006 | P83553 | Dermatopontin (Tyrosine-rich acidic matrix protein) (TRAMP) | |
| **LOC107330232** | 34614 | 61885 | Q01528; P83553 | Hemagglutinin/amebocyte aggregation factor (18K-LAF);Dermatopontin (Tyrosine-rich acidic matrix protein) (TRAMP) | |
| **LOC107330312** | 67432 | 69019 | | | |
| **LOC107330158** | 93005 | 95544 | | | |
| **LOC107330205** | 97052 | 103377 | Q01528 | Hemagglutinin/amebocyte aggregation factor (18K-LAF) | |
| **LOC107330284** | 108614 | 112414 | Q01528 | Hemagglutinin/amebocyte aggregation factor (18K-LAF) | |
| **LOC107330124** | 154755 | 156368 | Q8C4P0 | Uncharacterized protein KIAA1958 homolog | |
| **LOC107329866** | 187208 | 187804 | | | |
| | | | | **(SNP at 208400)** | |
| **LOC107330388** | 271722 | 293622 | P48960; Q24JV9 | CD97 antigen (Leukocyte antigen CD97) (CD antigen CD97) [Cleaved into: CD97 antigen subunit alpha; CD97 antigen subunit beta];Galectin-3-binding protein A (Lectin galactoside-binding soluble 3-binding protein A) | KOG4193:G- protein-coupled receptor |
| **LOC107330468** | 294995 | 301323 | Q19673 | Putative tyrosinase-like protein tyr-3 | |
| **LOC107330457** | 302642 | 304245 | | | |
| **LOC107329885** | 304354 | 312280 | Q9WTN0 | Geranylgeranyl pyrophosphate synthase (GGPP synthase) (GGPPSase) (EC 2.5.1.-) ((2E,6E)- | KOG0777:geranylgeranyl diphosphate synthase |

| | | | | | |
|---|---|---|---|---|---|
| | | | | farnesyl diphosphate synthase) (Dimethylallyltranstransferase) (EC 2.5.1.1) (Farnesyl diphosphate synthase) (Farnesyltranstransferase) (EC 2.5.1.29) (Geranylgeranyl diphosphate synthase) (Geranyltranstransferase) (EC 2.5.1.10) | |
| **LOC107329895** | 315833 | 327894 | Q9DBW0 | Cytochrome P450 4V2 (EC 1.14.14.-) (Docosahexaenoic acid omega-hydroxylase CYP4V2) (EC 1.14.14.79) | KOG0157:Cytochrome p450 |
| **LOC107329904** | 341135 | 341765 | Q8ITC7 | Neuropeptides capa receptor (Cap2b receptor) (Capability receptor) | KOG3656:receptor |
| **LOC107330438** | 344661 | 360273 | Q9DBW0 | Cytochrome P450 4V2 (EC 1.14.14.-) (Docosahexaenoic acid omega-hydroxylase CYP4V2) (EC 1.14.14.79) | KOG0157:Cytochrome p450 |
| **LOC107329913** | 360433 | 370933 | Q9DBW0 | Cytochrome P450 4V2 (EC 1.14.14.-) (Docosahexaenoic acid omega-hydroxylase CYP4V2) (EC 1.14.14.79) | KOG0157:Cytochrome p450 |
| **LOC107330484** | 372110 | 382570 | Q9DBW0 | Cytochrome P450 4V2 (EC 1.14.14.-) (Docosahexaenoic acid omega-hydroxylase CYP4V2) (EC 1.14.14.79) | KOG0157:Cytochrome p450 |
| **LOC107330491** | 385199 | 393983 | | | |
| **LOC107329921** | 446405 | 450470 | Q9UMS0 | NFU1 iron-sulfur cluster scaffold homolog, mitochondrial (HIRA-interacting protein 5) | KOG2358:NFU1 iron-sulfur cluster scaffold homolo |

**Supplementary Box 3. Genotype Environment Association GEA3.** Graph in a) shows the logistic model describing the presence (Probability=1) and absence (Probability=0) of the genetic variant as a function of the environmental gradient (DHW frequency). The frequency of the genotype occurrences in real data is represented by the grey bars. The map in b) show the average values of the environmental variable and the genotype frequency by sampling sites. Table in c) illustrates the annotations of the reference genome in the neighborhood (± 250 kb) of the significant SNP. For every locus, the table shows the starting and ending position on the scaffold, the best match of the sequence similarity search against the swissprot database (swissprot, E<$10^{-7}$), the annotation of the best match (annotation) and the related eukaryotic cluster of orthologs (KOGs).

a)

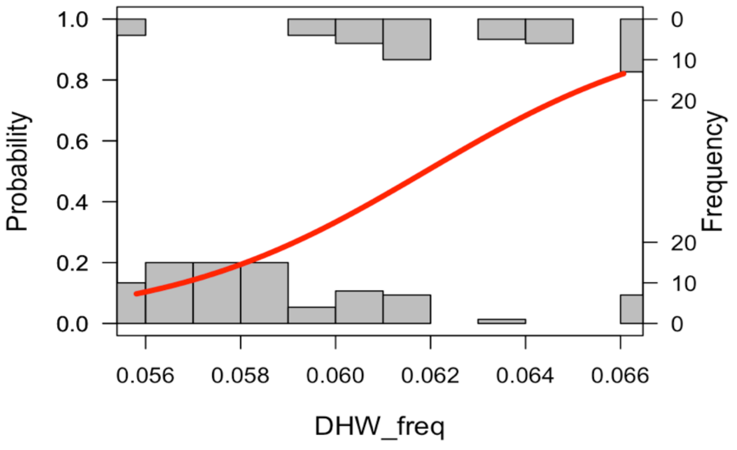


b)

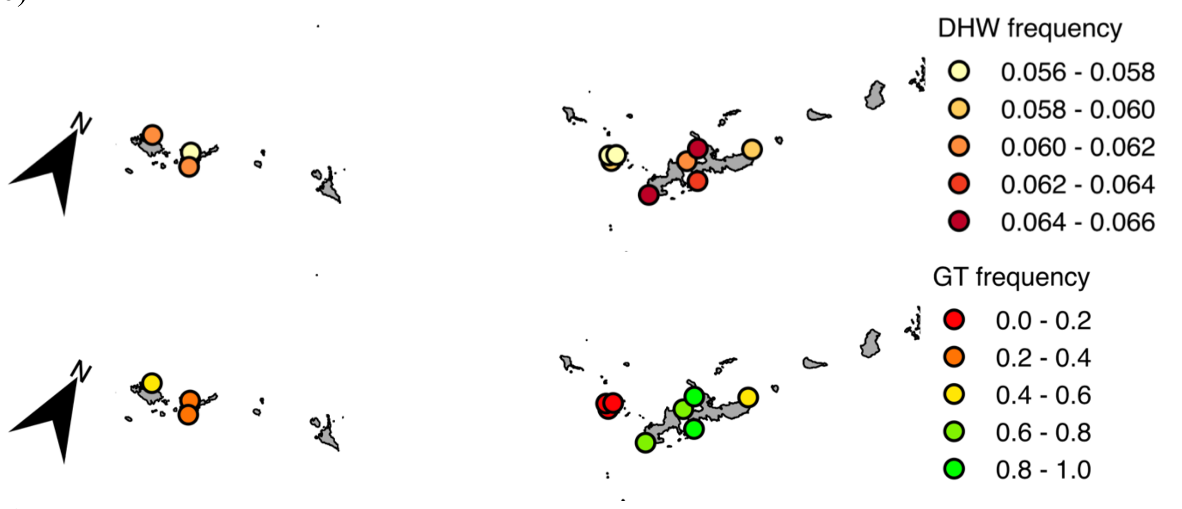


c)

| gene ID | position start | position end | swissprot | annotation | KOGs |
|---|---|---|---|---|---|
| **LOC107334806** | 452616 | 455315 | | | |
| **LOC107334807** | 455384 | 457531 | | | |
| **LOC107334825** | 463820 | 479184 | | | |
| **LOC107334822** | 481093 | 482278 | | | |
| **LOC107334808** | 497044 | 547778 | Q5S006 | Leucine-rich repeat serine/threonine-protein kinase 2 (EC 2.7.11.1) | KOG0192: protein kinase kinase kinase, KOG0619: leucine rich repeat |
| **LOC107334809** | 547671 | 548575 | | | |
| **LOC107334810** | 607102 | 607887 | | | |
| **LOC107334834** | 638325 | 654182 | | | |
| | | | **(SNP at 665651)** | | |
| **LOC107334811** | 712432 | 721913 | | | |
| **LOC107334835** | 850502 | 854864 | | | |
| **LOC107334814** | 862801 | 863746 | | | |
| **LOC107334815** | 881337 | 890355 | Q9P055 | JNK1/MAPK8-associated membrane protein (JKAMP) (JNK1-associated membrane protein) (JAMP) (Medulloblastoma antigen MU-MB-50.4) | KOG3744: jnk1 mapk8-associated membrane protein |
| **LOC107334838** | 893078 | 916936 | | | |
| **LOC107334837** | 897844 | 921305 | | | |

**Supplementary Box 4. Genotype Environment Association GEA4.** Graph in a) shows the logistic model describing the presence (Probability=1) and absence (Probability=0) of the genetic variant as a function of the environmental gradient (DHW frequency). The frequency of the genotype occurrences in real data is represented by the grey bars. The map in b) show the average values of the environmental variable and the genotype frequency by sampling sites. Table in c) illustrates the annotations of the reference genome in the neighborhood (± 250 kb) of the significant SNP. For every locus, the table shows the starting and ending position on the scaffold, the best match of the sequence similarity search against the swissprot database (swissprot, $E<10^{-7}$), the annotation of the best match (annotation) and the related eukaryotic cluster of orthologs (KOGs).

a)

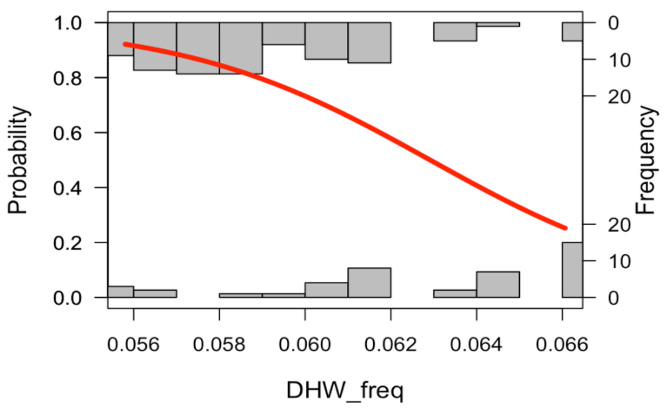


b)

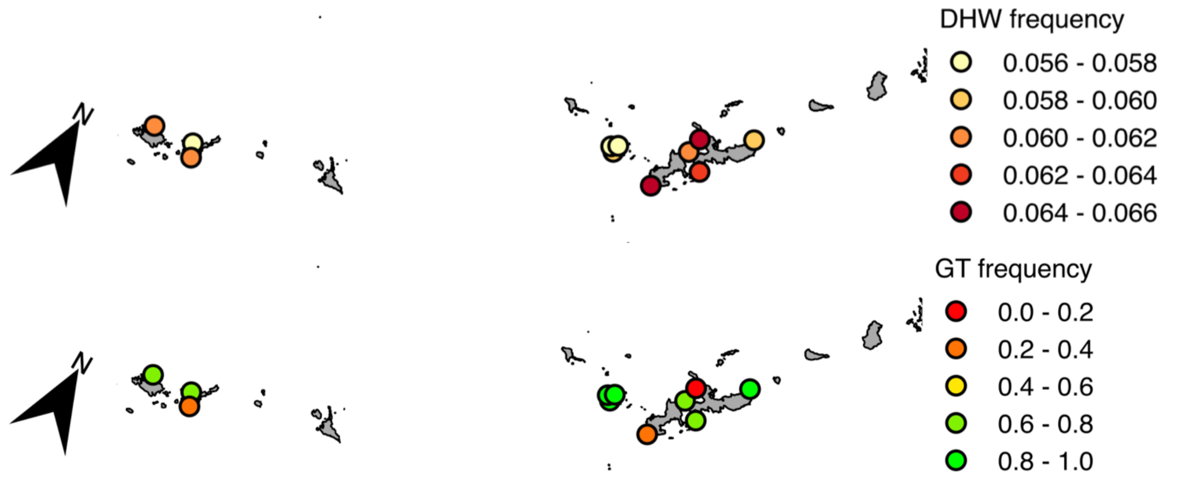


c)

| gene ID | position start | position end | swissprot | annotation | KOGs |
|---|---|---|---|---|---|
| | | | | (SNP at 27616) | |
| **LOC107344289** | 50930 | 51592 | | | |
| **LOC107344291** | 52950 | 59932 | | | |
| **LOC107344292** | 100498 | 101196 | | | |
| **LOC107344282** | 115535 | 116160 | | | |
| **LOC107344293** | 117963 | 119456 | Q96DM1 | PiggyBac transposable element-derived protein 4 | |
| **LOC107344294** | 141584 | 142972 | | | |
| **LOC107344295** | 144155 | 145962 | | | |
| **LOC107344296** | 148733 | 149463 | | | |
| **LOC107344307** | 199204 | 202345 | | | |
| **LOC107344283** | 203264 | 208238 | | | |
| **LOC107344298** | 265263 | 266592 | Q14191 | Werner syndrome ATP-dependent helicase (EC 3.6.4.12) (DNA helicase, RecQ-like type 3) (RecQ3) (Exonuclease WRN) (EC 3.1.-.-) (RecQ protein-like 2) | KOG0351: DNA helicase, KOG4373: Exonuclease 3'-5' domain containing 2 |

**Supplementary Box 5. Genotype Environment Association GEA5.** Graph in a) shows the logistic model describing the presence (Probability=1) and absence (Probability=0) of the genetic variant as a function of the environmental gradient (DHW frequency). The frequency of the genotype occurrences in real data is represented by the grey bars. The map in b) show the average values of the environmental variable and the genotype frequency by sampling sites. Table in c) illustrates the annotations of the reference genome in the neighborhood (± 250 kb) of the significant SNP. For every locus, the table shows the starting and ending position on the scaffold, the best match of the sequence similarity search against the swissprot database (swissprot, $E<10^{-7}$), the annotation of the best match (annotation) and the related eukaryotic cluster of orthologs (KOGs).

a)

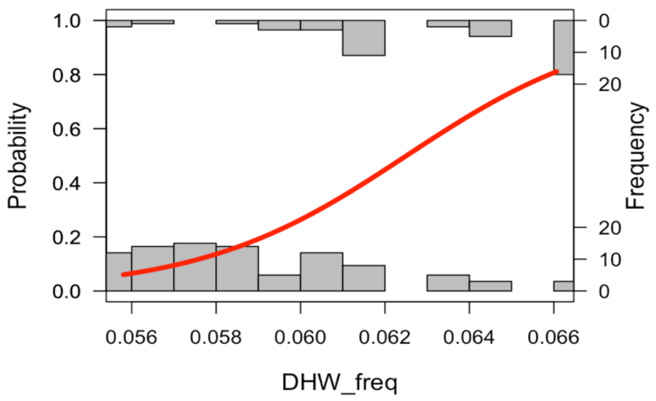


b)

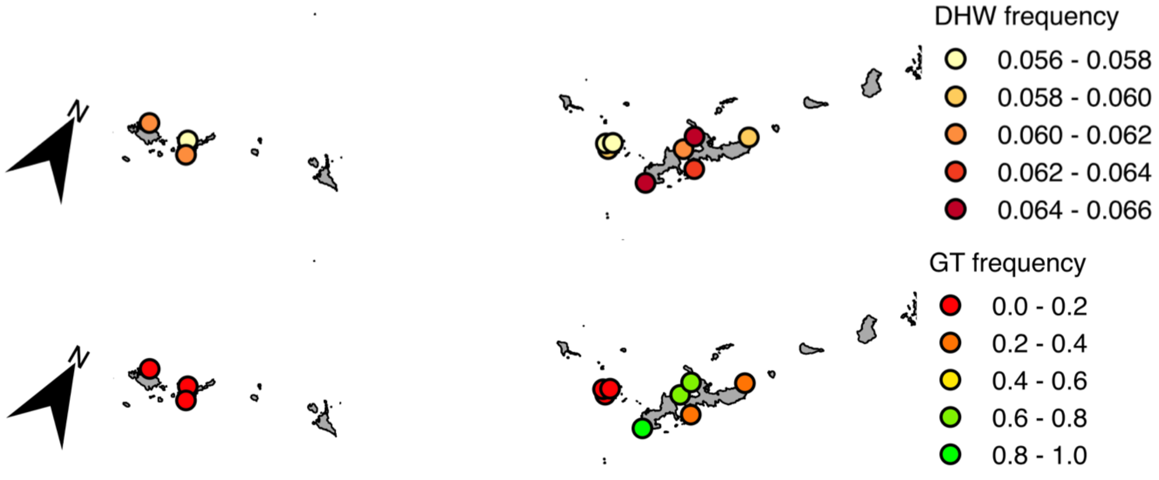


c)

| gene ID | position start | position end | swissprot | annotation | KOGs |
|---|---|---|---|---|---|
| **LOC107357078** | 2396 | 4597 | | | |
| | | | | (SNP at 16151) | |
| **LOC107357082** | 24194 | 26577 | | | |
| **LOC107357079** | 38606 | 41565 | | | |
| **LOC107357081** | 106905 | 160916 | O75665 | Oral-facial-digital syndrome 1 protein (Protein 71-7A) | |
| **LOC107357080** | 162832 | 163344 | Q14191 | Werner syndrome ATP-dependent helicase (EC 3.6.4.12) (DNA helicase, RecQ-like type 3) (RecQ3) (Exonuclease WRN) (EC 3.1.-.-) (RecQ protein-like 2) | KOG0351: DNA helicase, KOG4373: Exonuclease 3'-5' domain containing 2 |

**Supplementary Box 6. Genotype Environment Association GEA6.** Graph in a) shows the logistic model describing the presence (Probability=1) and absence (Probability=0) of the genetic variant as a function of the environmental gradient (DHW frequency). The frequency of the genotype occurrences in real data is represented by the grey bars. The map in b) show the average values of the environmental variable and the genotype frequency by sampling sites. Table in c) illustrates the annotations of the reference genome in the neighborhood (± 250 kb) of the significant SNP. For every locus, the table shows the starting and ending position on the scaffold, the best match of the sequence similarity search against the swissprot database (swissprot, E<$10^{-7}$), the annotation of the best match (annotation) and the related eukaryotic cluster of orthologs (KOGs).

a)

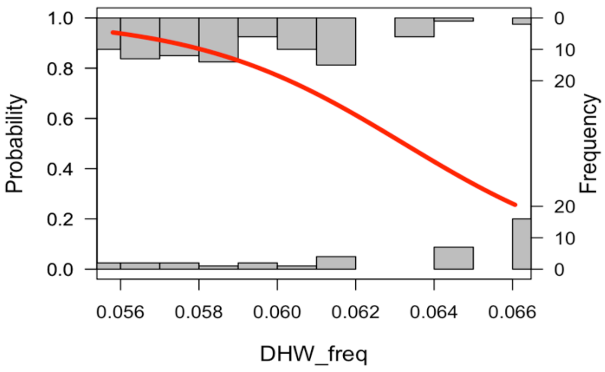


b)

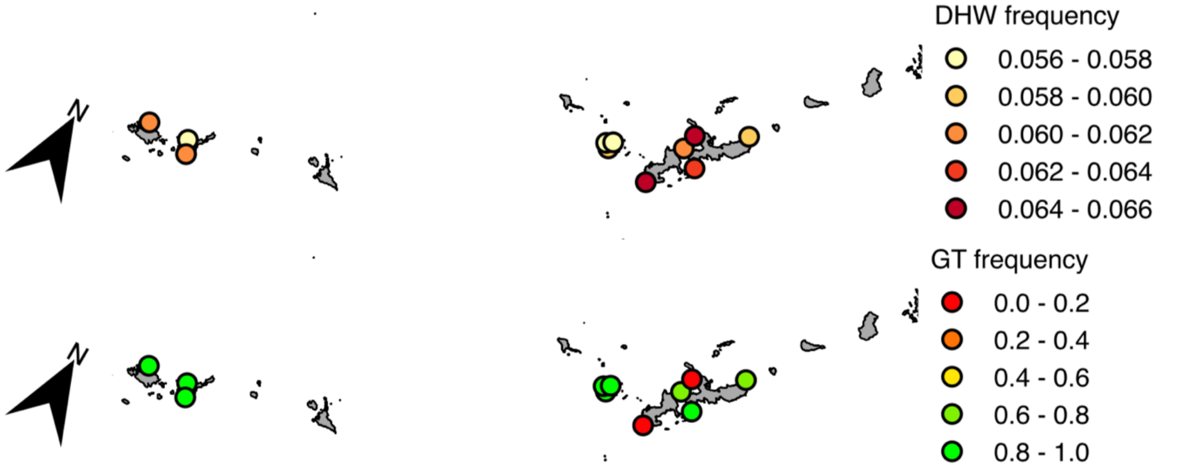


c)

| gene ID | position start | position end | swissprot | annotation | KOGs |
|---|---|---|---|---|---|
| **LOC107328703** | 5086 | 6804 | Q14191 | Werner syndrome ATP-dependent helicase (EC 3.6.4.12) (DNA helicase, RecQ-like type 3) (RecQ3) (Exonuclease WRN) (EC 3.1.-.-) (RecQ protein-like 2) | KOG0351: DNA helicase, KOG4373: Exonuclease 3'-5' domain containing 2 |
| | | | | **(SNP at 32233)** | |

**Supplementary Figure 4. Connectivity models.** The two models describe the relationship between genetic distance (Fst) and geographic distance calculated in two different ways. In a), geographic distance is sea distance and represents the dispersal costs calculated out of sea current data from the whole study period (1993-2010). In b), geographic distance is the aerial distance between sampling sites.

a)

Fst = -7.253e-04+(SD*1.551e-04)

p = 1.45e-09

Multiple R-squared = 0.7155

AIC = -234

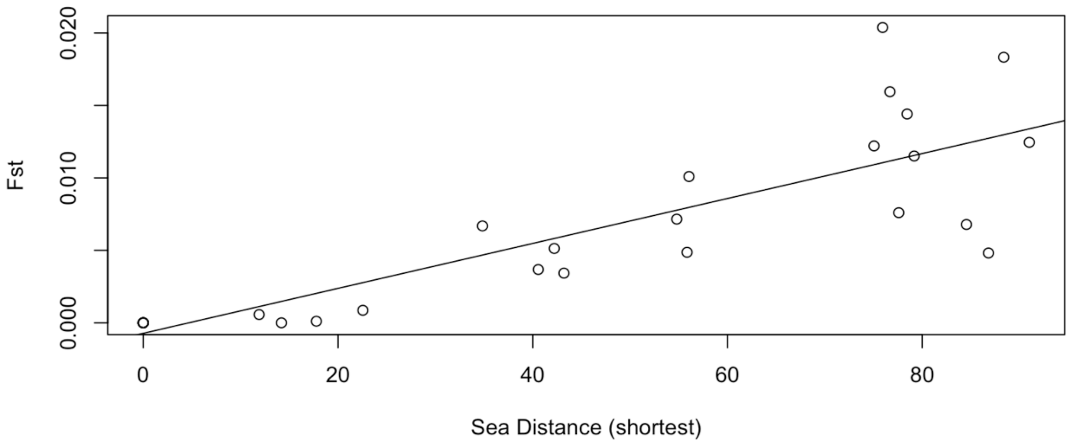


b)

Fst = -0.001051 +(SD*0.0001314)

p = 5.95e-07

Multiple R-squared = 0.62

AIC = -227

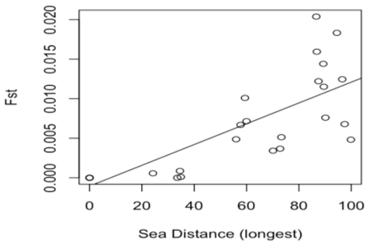


c)

Fst = -0.001049 +(SD*0.0001439)

p = 1.07e-07

Multiple R-squared = 0.66

AIC = -230

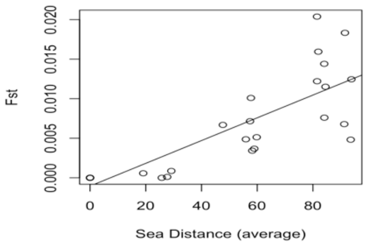


d)

Fst = 0.0016579+(AD*0.0027692)

p = 1.83e-07

Multiple R-squared = 0.66

AIC = -230

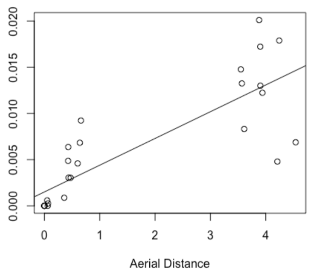

**Supplementary Figure 5. Example of pFst variation across study area.** The two maps show the pFst values connecting all the reefs of the study area to (a) and from (b) the same reef located in the center of Yaeyama islands (marked with an arrow).

a) pFst to Yaeyama

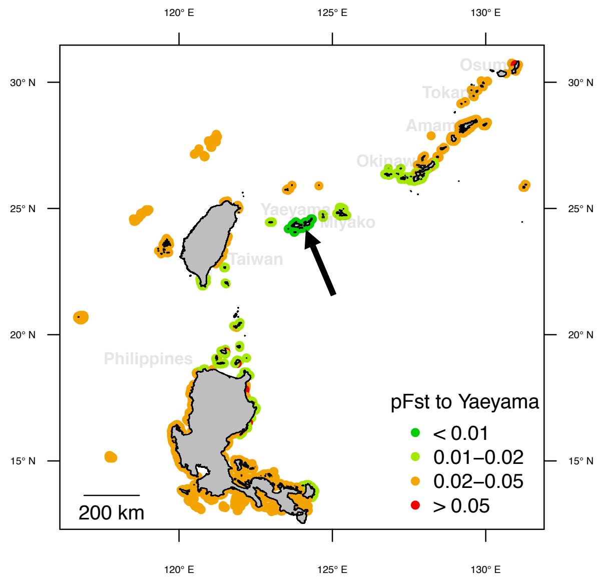


b) pFst from Yaeyama

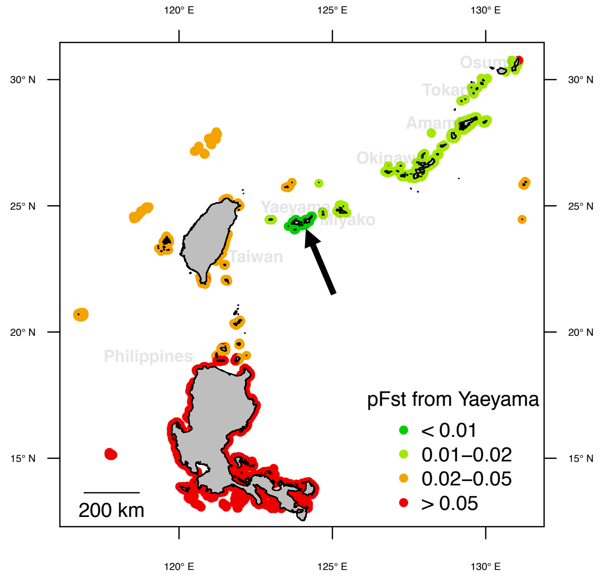

**Supplementary Figure 6. pFst from Northern Philippines.** The two maps show the pFst values connecting one reef in the north of Philippines (marked with an arrow) to all the reefs of the study area.

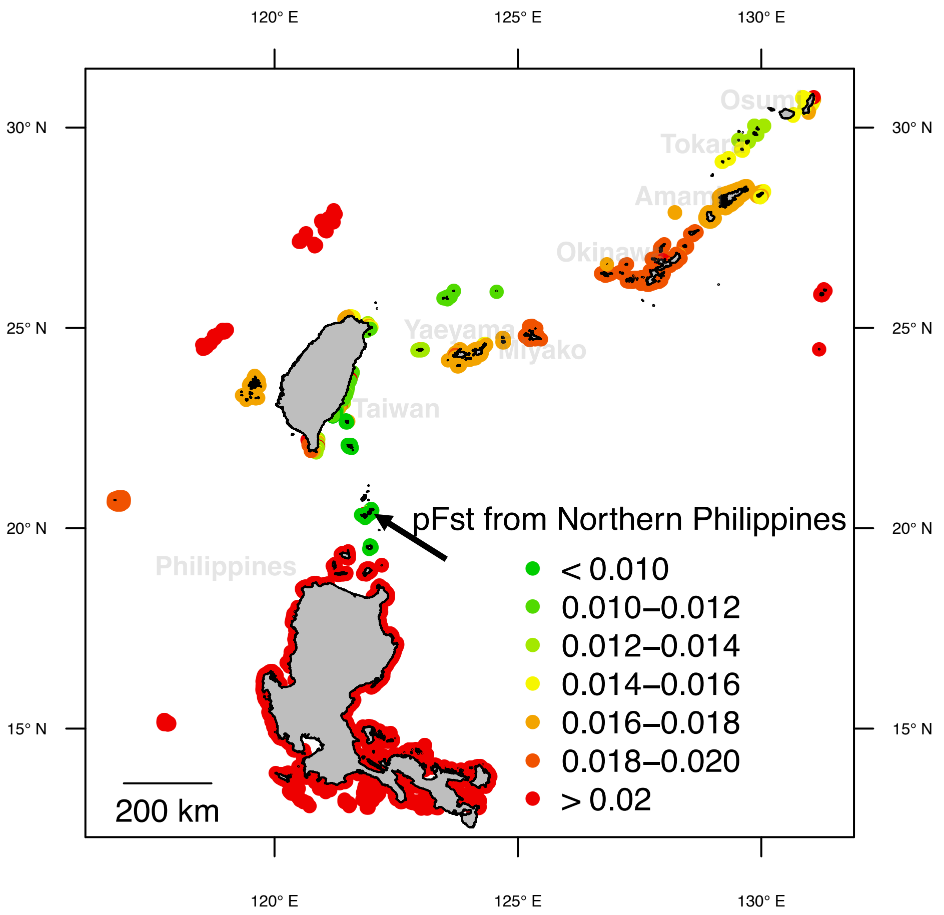
